# Nucleolar stress caused by arginine-rich peptides triggers a ribosomopathy and accelerates ageing in mice

**DOI:** 10.1101/2023.08.10.552792

**Authors:** Oleksandra Sirozh, Bomi Jung, Laura Sanchez-Burgos, Ivan Ventoso, Vanesa Lafarga, Oscar Fernández-Capetillo

**Author notes:** **Correspondence**: Oscar Fernandez-Capetillo or Vanesa Lafarga, Spanish National Cancer Research Centre (CNIO) Melchor Fernandez Almagro, 3 Madrid 28029, Spain, or.

## Abstract

Nucleolar stress (NS) has been associated to several age-related diseases such as cancer or neurodegeneration. To investigate the mechanisms of toxicity triggered by NS, we here used (PR)n arginine-rich peptides that are found in patients of some neurodegenerative diseases. Although these peptides accumulate at nucleoli and generate NS, how this translates into cellular toxicity is poorly understood. We here reveal that whereas (PR)n expression leads to an overall decrease in protein abundance, this occurs concomitant with an accumulation of free ribosomal (r) proteins in the cytoplasm, a hallmark of ribosomopathies. Conversely, cells with acquired resistance to (PR)n peptides present global downregulation of r-proteins and low levels of mTOR signaling. In mice, systemic expression of (PR)_97_ drives widespread NS and accelerated ageing, which is associated to an increased expression of r-proteins and mTOR hyperactivation. Furthermore, the reduced lifespan of (PR)_97_-expressing mice was alleviated by the mTOR inhibitor rapamycin. Importantly, we show that the generalised accumulation of free r-proteins is a common outcome in response to chemical or genetic perturbations that trigger NS, such as Actinomycin D, TIF-IA depletion, or the expression of mutant HMGB1 variants recently associated to rare human diseases. Together, our study presents *in vivo* evidence supporting the role of NS as a driver of ageing, and provides a general framework to explain the toxicity caused by NS in mammalian cells.

## Introduction

Nucleoli are the largest membrane-less nuclear organelles, best known for their central role in ribosome biogenesis^1,2^. Nevertheless, proteomic studies revealed that nucleoli harbour hundreds of proteins involved in multiple additional functions^3^ such as DNA repair and recombination, telomere maintenance or stress responses (reviewed in ^4^). The nucleolus is primarily formed by phase separation, driven by low-affinity interactions between ribosomal DNA (rDNA) and/or ribosomal RNA (rRNA) with nucleolar factors. The accumulation of proteins at nucleoli is mediated by nucleolar localization sequences, from which more than 50% of the amino acids are positively charged lysines and arginines^5,6^. First noted in 1985^7^, nucleolar proteins often change their localization upon various forms of stress, and this redistribution is often used as indicative of nucleolar stress (NS). Today, NS collectively refers to a wide range of chemical or genetic perturbations that target nucleoli and alter their shape and function^8,9^. Examples of NS-inducers include RNA polymerase I inhibitors^7^, viral proteins^10^, UV-light^11^, heat shock^12^ and DNA-damaging drugs^13,14^. Regardless of the insult, NS triggers the activation of P53, even in the absence of DNA damage^15^. Importantly, numerous data indicate that while P53 contributes to NS-driven toxicity, this stress ultimately kills cells by P53-independent mechanisms that remain to be deciphered^16^.

In addition to inducing NS with external agents, nucleolar alterations have been frequently documented in human disease^17^. For instance, aberrations in nucleolar shape and number are associated to poor prognosis in several cancer types^18^, and mutations or altered expression of r-proteins are often found in cancer cells^19,20^. Furthermore, strategies targeting nucleoli are being investigated as anti-cancer therapies^21^. Besides cancer, NS has been particularly associated to neurodegenerative diseases, perhaps due to the higher translational demand of neurons than that of other cell types^22^. In this regard, NS-inducing arginine-rich dipeptide repeats (DPRs), such as poly(PR) or poly(GR), have been found in patients of amyotrophic lateral sclerosis and spinocerebellar ataxia type 36, driven by mutations in *C9ORF72* and *SCA36*, respectively^23^. These DPRs accumulate at nucleoli, disrupt nucleolar function and lead to cell death^24^. Yet again, a full understanding of the mechanism of toxicity of these DPRs is still lacking.

Finally, and despite not being included as one of the “hallmarks of ageing”^25^, numerous evidences indicate a role for nucleoli in ageing^26^. Early studies in yeast revealed that rDNA instability accumulated during an organism’s lifespan and already suggested that safeguarding nucleolar function could be a life-extending strategy^27^. Accordingly, some of the best-known modulators of survival, such as the mTOR kinase, are regulators of ribosome biogenesis and r-protein synthesis ^28,29^. More recently, nucleolar size was shown to be inversely correlated with lifespan in several organisms, and depletion of the nucleolar factor fibrillarin shrank nucleoli and extended survival in C elegans^30^. However, genetic proof showing that NS accelerates ageing is still missing. Here, using (PR)n expression as a tool to induce NS, we set out to investigate the mechanisms of NS-toxicity and its impact on mammalian lifespan.

## Results

### (PR)_97_ expression drives the accumulation of free cytoplasmic r-proteins

To investigate the mechanisms mediating the toxicity of arginine-rich peptides, we generated a human osteosarcoma U2OS cell line that enabled doxycycline (dox)-inducible expression an HA-tagged (PR)_97_ polypeptide (U2OS(PR)^97^). We chose U2OS as this cell line was originally used by the McKnight laboratory to illustrate the toxicity of (PR)n and (GR)n peptides^24^. Consistent with their study, (PR)_97_ peptides accumulated at nucleoli, triggered NS (reflected as an increase in the nucleolar area) and killed cells (**Fig. 1A-C**). Moreover, drugs known to generate NS such as actinomycin D (ActD) or the RNA Pol I inhibitor CX-541, did not increase cell death in dox-treated U2OS(PR)^97^ cells (**Fig. S1**), supporting that NS is the main insult mediating the toxicity of (PR)n peptides. Of note, previous studies indicate that cell death triggered by arginine-rich peptides is at least partly mediated by P53^31,32^. However, while we detected a slight induction of P53 in response to a prolonged exposure to (PR)n peptides, synthetic (PR)_20_ peptides killed P53-proficient and deficient HCT-116 cells similarly, and CRISPR-mediated deletion of P53 did not rescue the viability of U2OS cells expressing (PR)_97_ (**Fig. S2**). These observations are consistent with decades of work, which have indicated that whereas P53 plays a role in the response to NS, this insult ultimately kills cells in a p53-independent manner that remains to be understood^16,17^.

**Figure 1.**
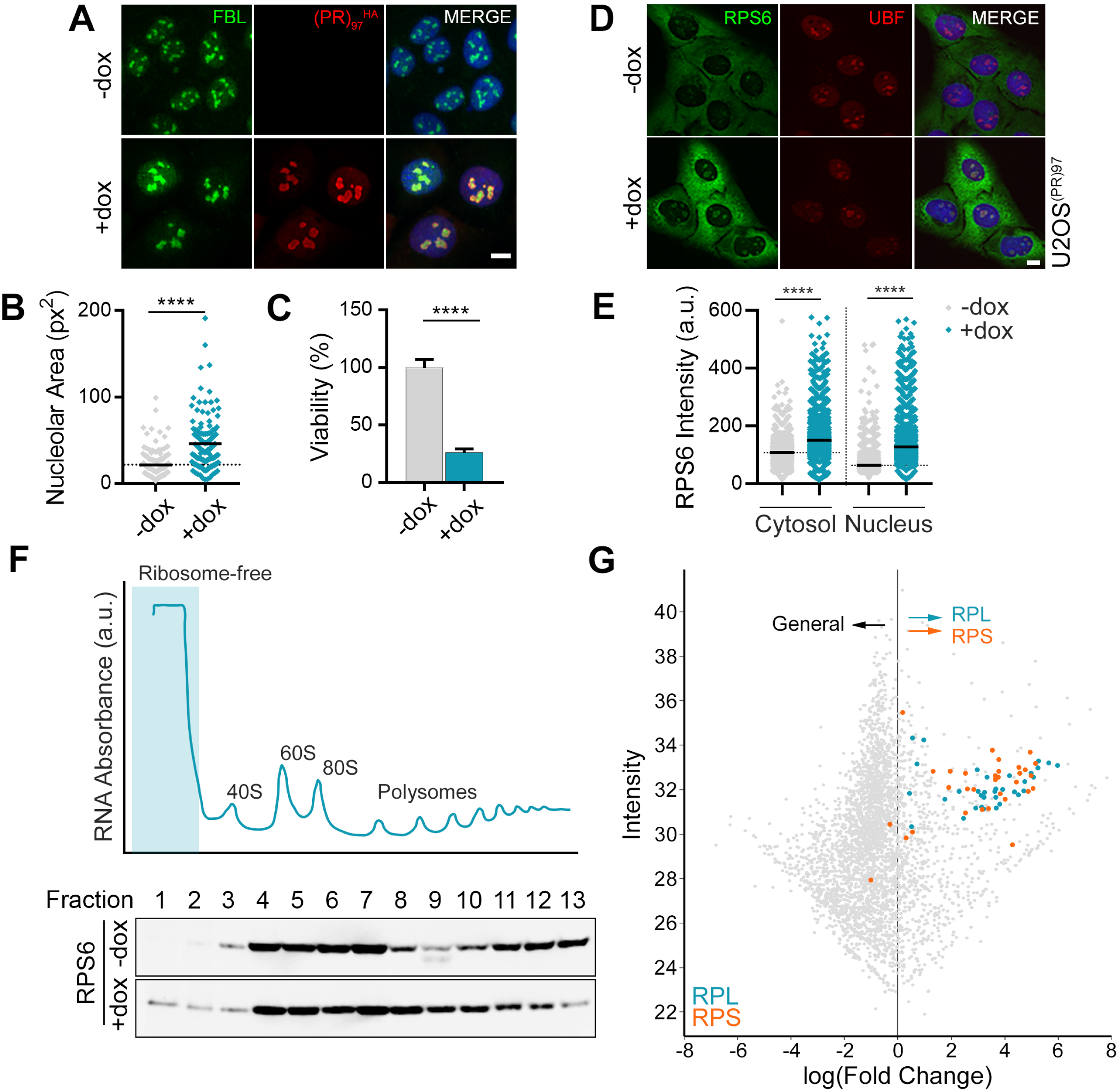
(PR)_97_ peptides trigger NS and the accumulation of free r-proteins. (A) Representative immunofluorescence of the nucleolar factor FBL (green) and (PR) ^HA^ (red), in U2OS(PR)^97^ cells following 48 hrs of treatment with dox. Scale bar (white), represents 10 μm. (B) HTM-mediated quantification of nucleolar area, as quantified by FBL staining, from the images provided in (A). Black lines indicate mean values. (C) Viability by HTM-mediated quantification of nuclei, stained by DAPI, after 48 hrs of treatment with dox. (D) Representative immunofluorescence of RPS6 (green) in dox treated U2OS(PR)^97^ cells. UBF (red) was used to stain nucleoli. (D) HTM-mediated quantification data shown in (D). Black lines indicate mean values. (F) Sucrose gradient fractionation of protein extracts from U2OS^PR97^ cells, treated with dox for 48h. RNA absorbance indicates the distribution of RNA between the different fractions. WB of the distribution of RPS6 between the different fractions. Note that RPS6 is detected in the ribosome-free fractions (1 and 2) of dox-treated cells, but not untreated cells. (G) Intensity and fold-change of all proteins identified by proteomic analysis of the ribosome-free fraction of dox-treated (48 hrs) U2OS(PR)^97^ cells compared to untreated cells. r-proteins are marked with orange (40S) and blue (60S subunit). Data information: \*\*\*\**P* < 0.0001; *t*-test.

To investigate the mechanisms driving (PR)n toxicity, we focused on the effects of these peptides on RNA metabolism. In this regard, we recently showed that due to their high affinity for RNA, these peptides decorate all cellular RNAs leading to a generalized displacement of RNA-binding proteins (RBPs) from mRNA^33^. Based on these findings, and given that (PR)n peptides primarily accumulate at nucleoli, and that the majority of cellular RNA is rRNA, we hypothesized that these peptides should have a preferential impact on mature rRNA biogenesis, thereby limiting the availability of processed rRNA molecules and consequently triggering an accumulation of free r-proteins in the cytoplasm. Consistent with this view, High-Throughput Microscopy (HTM)-dependent quantification revealed the accumulation of RPS6 in dox-treated U2OS(PR)^97^ cells (**Fig. 1D-E**). To evaluate whether the accumulation of RPS6 reflected an increase in free r-proteins, we conducted sucrose gradient fractionations. These experiments confirmed the accumulation of RPS6 in the ribosome-free fraction in response to (PR)_97_ expression (**Fig. 1F**). Consistent with the general downregulation of translation that has been repeatedly documented as a response to arginine-rich peptides^33–35^, proteomic analyses of the ribosome-free fraction revealed a generalized downregulation of protein levels in dox-treated U2OS(PR)^97^ cells (**Fig. 1G**). In striking contrast, (PR)_97_ expression led to a collective increase in the levels of free r-proteins (**Fig. 1G**). Of note, proteomic analysis of the polysome fraction revealed that there was a specific decrease of RPLs on polysomes, which we previously demonstrated was due to the inability of the 60S particle to bind to 40S-bound polysomes at initiation sites on (PR)n-decorated mRNA^33^ (**Fig. S3**). Together, these experiments show that the toxicity caused by (PR)n peptides is associated to a generalised increase of free r-proteins in the cytoplasm, which is a hallmark of ribosomopathies.

### Resistance to NS correlates with low levels of ribosome biogenesis

To further explore the mechanisms of toxicity of arginine-rich peptides, we generated two independent clones of mouse NSC34 motor neuron-like cells that developed adaptative resistance to synthetic (PR)_20_ upon continuous culture in the presence of these peptides (NSC34^R^). Isolated clones were expanded and showed significant resistance when exposed to increasing doses of (PR)_20_ peptides (**Fig. 2A**). Moreover, whereas NSC34^R^ cells were not resistant to other drugs that activate the P53-dependent response such as cisplatin, camptothecin or inhibitors of the ATR kinase, they presented resistance to several treatments that induce NS, such as synthetic (PR)_20_ peptides, ActD or CX-541 (**Fig. 2B**). These results indicate that NSC34^R^ cells present a generalised resistance to NS, thus becoming a useful model to investigate the mechanisms driving NS-dependent toxicity in mammalian cells.

**Figure 2.**
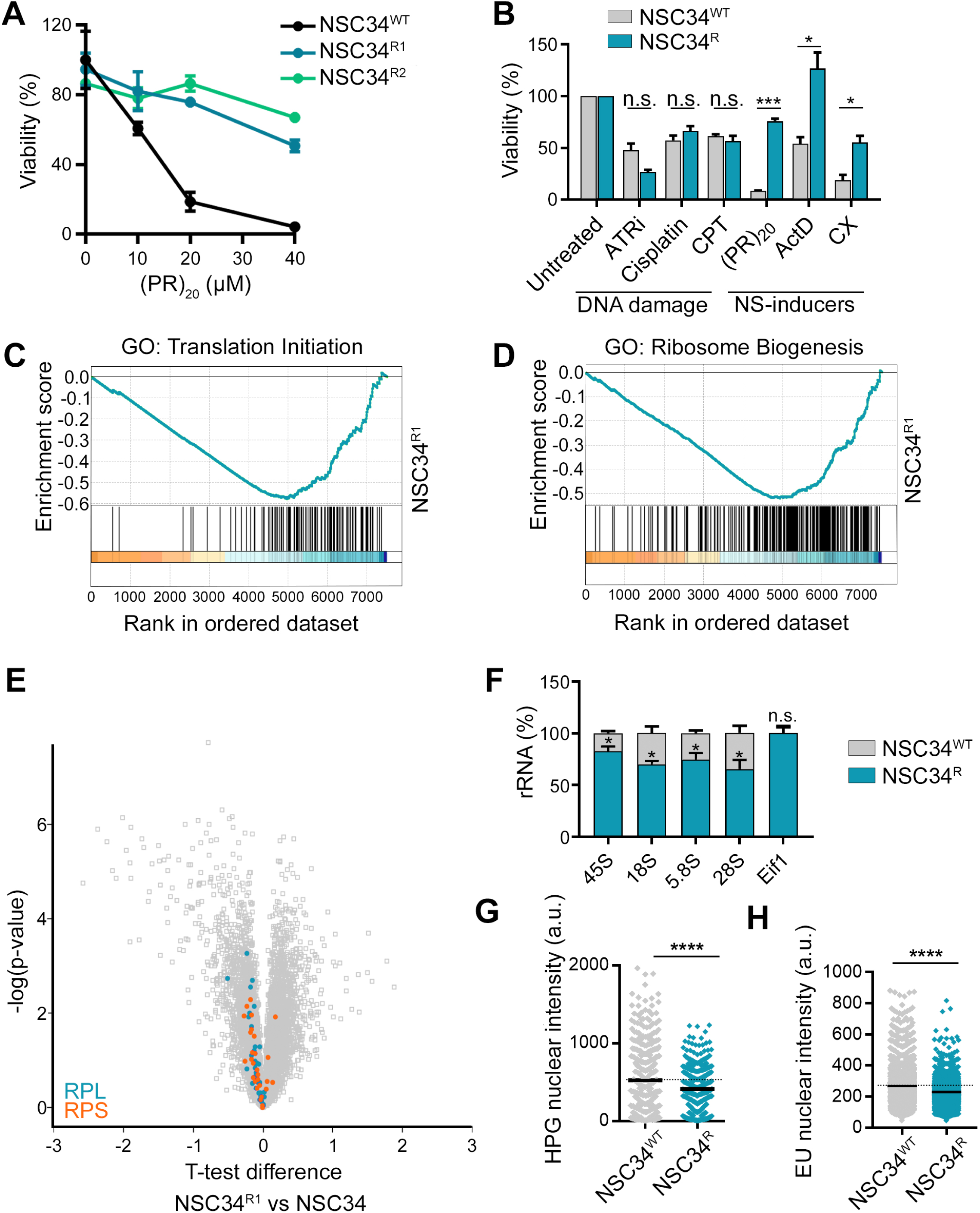
Reduced nucleolar activity in (PR)n-resistant NSC34 cells. (A) Normalised viability, as evaluated with a CellTiter-Glo luminescent assay, of wild-type NSC-34 cells (black), and two spontaneously resistant clones (NSC34^R1^ (blue) and NSC34^R^^2^ (green)), treated with increasing doses of (PR)_20_ peptides for 24 hrs. (B) Normalized viability, as evaluated with CellTiter-Glo luminescent assay, of NSC34^WT^ (grey) and NSC34^R1^ (blue), treated with several genotoxic compounds (ATRi, 20 μM; cisplatin, 25 μM; and camptothecin (CPT), 8 μM) and NS-inducing treatments ((PR)_20,_ 20 μM; ActD, 5 nM and CX-5461, 10 nM), for 24 hrs. (C,D) GSEA enrichment plots for the gene ontology classes corresponding to ribosome biogenesis (C), and translation initiation (D) derived from RNA-Seq data comparing NSC34^R1^ cells with the parental NSC34 cell line. (E) Volcano plot of all proteins identified by proteomic comparison of NSC34^R1^ cells with the parental NSC34 cell line. Ribosomal proteins are marked with orange (40S subunit) and blue (60S subunit). (F) rRNA precursor transcript (45S) and processed rRNA (18S, 5.8S, and 28S) levels in wild-type NSC34^WT^ cells (grey) and spontaneously resistant NSC34^R^ (blue), quantified by qRT-PCR, and normalised against *Eif1* mRNA levels. (G, H) HTM-mediated quantification of the HPG levels per nucleus (G) and EU levels per cell (H) in NSC34^WT^ and spontaneously resistant NSC34^R^ cells. EU and HPG were added 30 minutes prior to fixation. Black lines indicate mean values. Data information: n.s.: non-significant, \**P* < 0.05; \*\*\**P* < 0.001; \*\*\*\**P* < 0.0001 *t*-test.

To investigate the molecular features associated with NS resistance in NSC34^R^ cells, we compared their proteomes to that of parental NSC34 cells. Gene ontology analyses revealed all aspects related to ribosome biogenesis and translation as those most significantly downregulated in the resistant cells (**Fig. 2C,D** and **S4A**). In agreement with this findings, NSC34^R^ cells presented a global decrease in total r-protein levels (**Fig. 2E**). Importantly, these observations were confirmed in an independent clone of NS-resistant NSC34 cells (**Fig. S4B-D)**. Concomitant to these changes in r-protein levels, NSC34^R^ cells also had lower levels of rRNA synthesis, measured by HTM-quantification of 5-Ethynyl Uridine (EU) incorporation, or by quantitative RT-PCR of specific rRNA transcripts (**Fig. 2F,G**). Consistent with these findings, translation levels, quantified by the incorporation of L-Homopropargylglycine (HPG) using HTM, were lower in NSC34^R^ cells (**Fig. 2H**). In summary, resistance to NS correlates with a general downregulation of ribosome biogenesis, which is opposite to the accumulation of free r-proteins that is observed upon exposure to (PR)n peptides.

### Transcriptomic analyses identify mTOR inhibitors as drugs counteracting (PR)n toxicity

We next tried to identify drugs that can trigger transcriptional changes equivalent to those found in NS-resistant cells. To identify such compounds, we used proteomic expression data from NSC34^R^ cells to interrogate the Connectivity Map (CMap) database (https://clue.io/). This web server stores the transcriptional signatures of more than 5,000 small molecules and integrates tools that can be used to search for similarities to a user-provided signature^36^. Drug Set Enrichment Analyses (DSEA), based on the connectivity data obtained from CMap, identified a significant enrichment of mTOR inhibitors among drugs with a signature that resembles that of NSC34^R^ cells (**Fig. 3A, B**). In agreement with these findings, Western Blotting (WB) revealed reduced levels of S6K1 or RPS6 phosphorylation in NSC34^R^ cells, indicative of low mTOR activity (**Fig. 3C**). Based on these observations, we next tested if mTOR inhibition could alleviate the toxicity of (PR)_n_ peptides. To do so, we generated a dox-inducible (PR)_97_ expression system in NSC34 cells, analogous to the previously described one in U2OS cells (NSC34(PR)^97^). In this cell model, the use of rapamycin (Rapa) or the ATP competitive mTOR inhibitor Torin-1 significantly rescued the toxicity driven by (PR)_97_ expression (**Fig. 3D, E**). Rapa treatment also rescued the neurite atrophy observed in dox-treated NSC34(PR)^97^ cells that were previously differentiated into neurons (**Fig. 3D, F**). In contrast to the rescue of (PR)n toxicity that is seen upon mTOR inhibition, mouse embryonic fibroblasts (MEF) deficient in TSC complex 2 (TSC2), and thus presenting constitutively elevated mTOR activity, were hypersensitive to synthetic (PR)_20_ peptides (**Fig. 3G**). Taken together, these results indicate that the toxicity of (PR)n peptides can be alleviated by mTOR inhibition, which recapitulates the phenotype of NSC34^R^ cells. Noteworthy, CMap analyses also identified inhibitors of IGF-1, AKT or PI3K as potentially counteracting (PR)n toxicity, all of which, like mTOR, having been investigated as regulators of ribosome biogenesis and ageing (**Fig. S5**).

**Figure 3.**
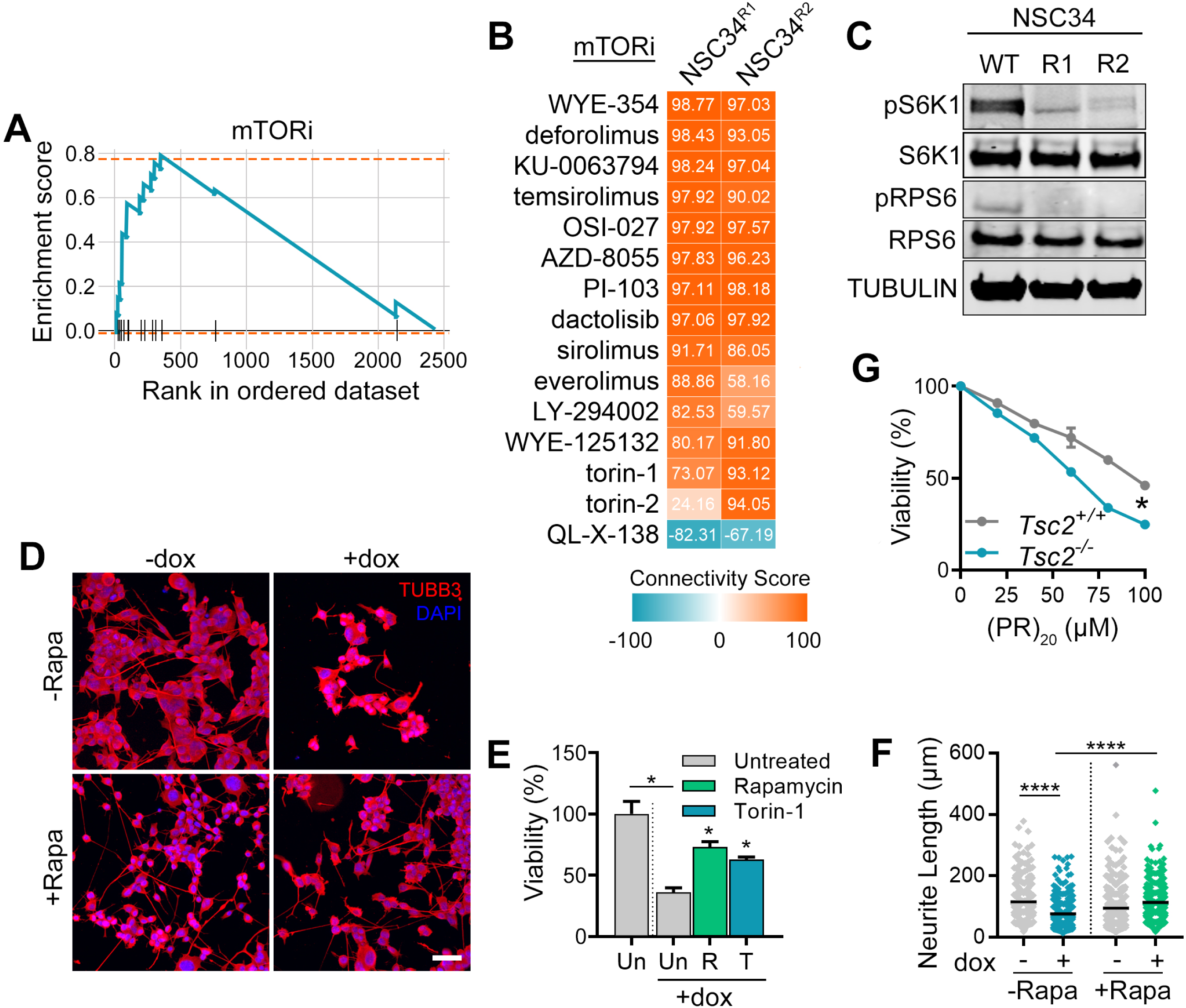
mTOR inhibition recapitulates the (PR)n-resistance of NSC34^R^ cells. (A) Drug set enrichment analysis (DSEA) for the most significantly enriched drug class (mTOR inhibitors), that trigger a transcriptional signature mimicking that found in NSC34^R1^ and NSC34^R2^ cells. (B) CMap connectivity scores (CS) of the mTOR inhibitor compound set, illustrating the similarity of their transcriptional signature to that of NSC34^R1^ and NSC34^R2^ cell. The CS is represented by the colour gradient, and the value indicated in each cell. (C) WB evaluating the levels of r-proteins and mTOR signalling in wild-type and (PR)n-resistant NSC34 cells. Total and phosphorylated levels of S6K1 and RPS6 are shown, along with TUBULIN as a loading control. (D) Immunofluorescence of the neuronal marker TUBB3 (red) in differentiated NSC-34(PR)^97^ cells treated with dox for 48 hrs, with or without 50 nM rapamycin. Nuclei were stained with DAPI (blue). Scale bar (white), 10 μm. (E) Average normalised viability, as evaluated with manual counting of live cells, of two independent inducible NSC-34(PR)^97^ cell lines, with and without dox, treated with rapamycin (50 nM) or torin-1 (50 nM) over 48 hrs. (F) Quantification of the neurite lengths from the experiment shown in (D). (G) Normalised viability, as evaluated with a CellTiter-Glo luminescent assay, of *Tsc2* WT or knockout MEF, treated with increasing doses of (PR)_20_ peptide for 24 hrs (n=2). Data information: * *P* < 0.05; **** *P* < 0.0001; *t*-test.

### Systemic expression of (PR)_97_ peptides accelerates ageing in mice

To investigate the consequences of NS in an animal model, we generated mice that enabled a generalised expression of (PR)_n_ peptides (PR knock-in mice; PR^KI^). To do so, we used a previously developed strategy whereby expression of the reverse tet-controlled transactivator (rtTA) is driven by the *Rosa26* promoter, and the transgene of interest is placed under the control of tet operator sequences (*tetO*) is inserted at the *Col1a1* locus^37^. This split system reduces the leakiness of tetracycline-regulated expression systems, which is particularly relevant for toxic transgenes. The transgene was a codon-optimised (PR)_97_ sequence, avoiding repetitive sequences, followed by a human influenza hemagglutinin (HA) tag (**Fig. S6A**). Administration of dox in the drinking water to 5-week-old PR^KI/KI^ mice led to the expression of (PR)_97_^HA^ in multiple organs, such as the intestines, skin, pancreas liver and bone marrow (**Fig. 4A** and **S6B**). Immunohistochemistry (IHC) against the HA-tag showed cytoplasmic and nuclear expression of the (PR)_97_^HA^ peptide, with examples of nucleolar accumulation also detected (**Fig. 4A**). The nucleolar accumulation of (PR)_97_^HA^, as seen *in vitro*, was particularly evident when the peptide was detected by immunofluorescence (IF) (**Fig. 4B**). Systemic expression of (PR) ^HA^ led to a significant reduction in mouse lifespan in a transgene dose-dependent manner (**Fig. 4C**). Interestingly, the cause of death in PR^KI/KI^ mice was the onset of a progeroid phenotype, as evidenced by multiple ageing-associated phenotypes, such as the appearance of cataracts, hair greying, kyphosis, loss of body weight, thinning of the skin, and the replacement of bone marrow hematopoietic cells with adipose tissue (**Fig. 4D-G** and **Fig. S6C-F**). Thus, systemic expression of (PR)n peptides drives a generalized accumulation of NS and accelerates ageing in mice.

**Figure 4.**
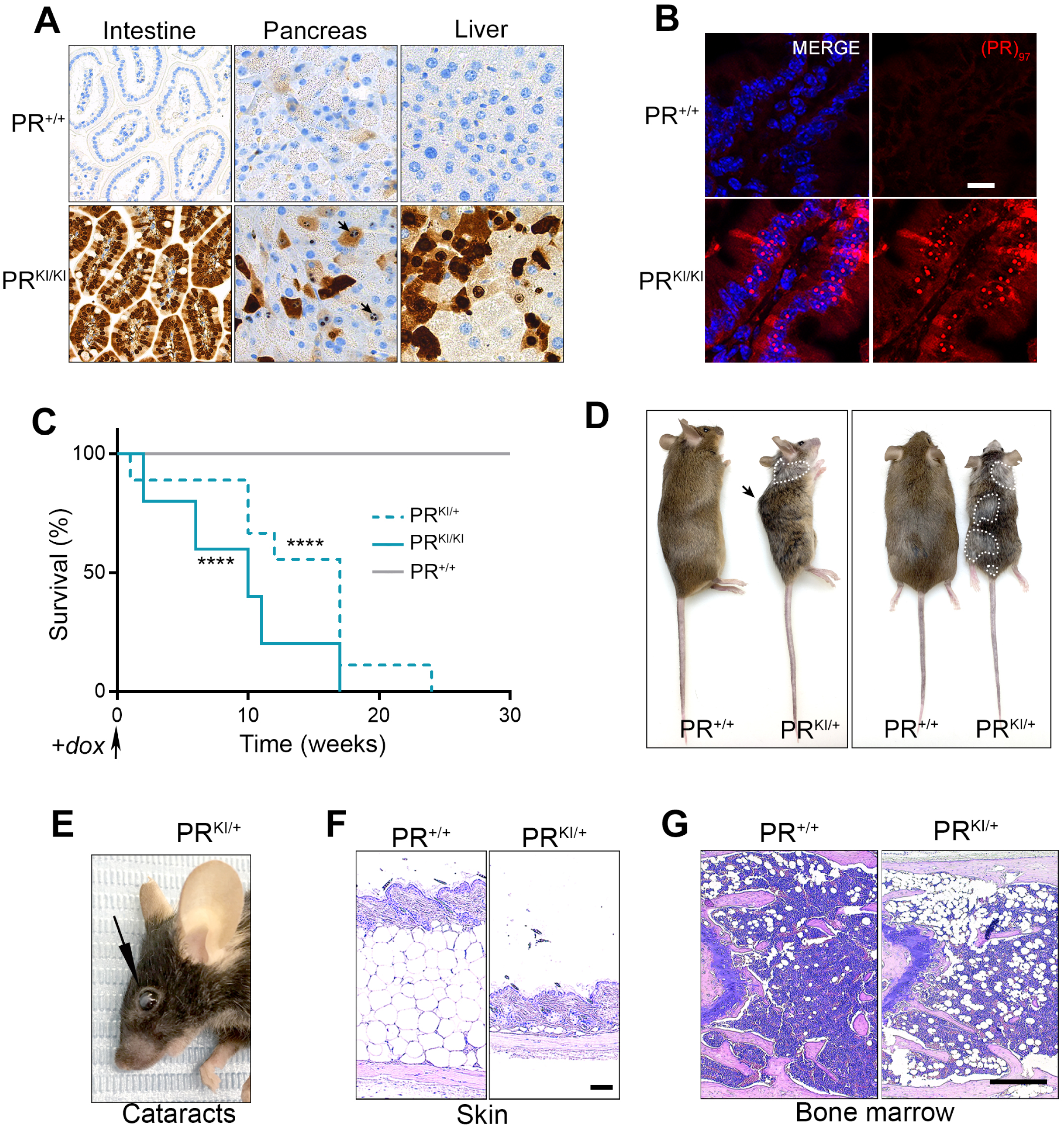
Systemic (PR) ^HA^ expression accelerates ageing in mice. (A) HA IHC in sections of intestine, pancreas, and liver from littermate PR^+/+^ and PR^KI/KI^ mice, 3 weeks after treatment with dox in the drinking water. Black arrows indicate examples of nucleolar (PR) ^HA^ localisation in pancreatic cells. (B) IF of (PR) ^HA^ peptides (red) in pancreas sections from littermate PR^+/+^ and PR^KI/KI^ mice, 3 weeks after treatment with dox in the drinking water. (C) Kaplan-Meyer plot representing the lifespan of non-inducible (PR^+/+^; *n* = 8), heterozygous inducible (PR^KI/+^, *n* = 9), and homozygous inducible (PR^KI/KI^, *n* = 5) mice, after starting dox treatment at 5 weeks. (**** *P* < 0.001; Mantel-Cox test). (D) Representative lateral and dorsal images of non-inducible PR^+/+^ and inducible PR^KI/+^ littermates after 20 weeks of treatment with dox. The black arrow indicates the presence of kyphosis, and dotted white lines delimit areas of grey fur in PR^KI/KI^ mice. (E) Representative image of a cataract (black arrow) in an inducible PR^KI/KI^ mouse after 20 weeks of treatment with dox. (F, G) Haematoxylin and eosin (H&E) staining of skin (F) and femoral bone marrow (G) from PR^+/+^ and PR^KI/+^ littermates after 20 weeks of treatment with dox to illustrate the decrease in skin thickness and the replacement of hematopoietic cells with adipose tissue in (PR) ^HA^ expressing mice. Scale bars represents 100 μM (F) and 500 μm (G).

### Ageing of (PR)_97_-expressing mice is associated to NS, mTOR hyperactivation and accumulation of ribosomal proteins

Next, we evaluated if our molecular observations from analysing the response to (PR)n peptides *in vitro*, were also recapitulated in mice. In PR^KI/KI^ MEF, dox treatment led to an accumulation of (PR)_97_^HA^ peptides at nucleoli, which led to NS, as evidenced by an increase in nucleolar area and a reduction in translation and cell viability (**Fig. S7**). Similarly, a 3-week treatment with dox led to NS in PR^KI/KI^ mice, detected as an increase in the area of nucleolar fibrillarin (FBL) in several organs, such as the liver (**Fig. 5A,B**) or intestine (**Fig. S8A,B**). To investigate the alterations in gene expression associated with the progeroid phenotype triggered by (PR)_97_^HA^ peptides *in vivo*, we performed RNA-seq on livers of PR^+/+^ and PR^+/KI^ mice treated with dox for 17 weeks. Gene Set Enrichment Analyses (GSEA) revealed a significant enrichment in the category of ‘cytoplasmic r-proteins’ in the liver of PR^+/KI^ mice (**Fig. 5C**). Consistently, there was a generalized increase in the expression of r-protein-coding genes in the liver of dox-treated PR^+/KI^ mice (**Fig. 5D**), as confirmed by WB (**Fig. 5E**).

**Figure 5.**
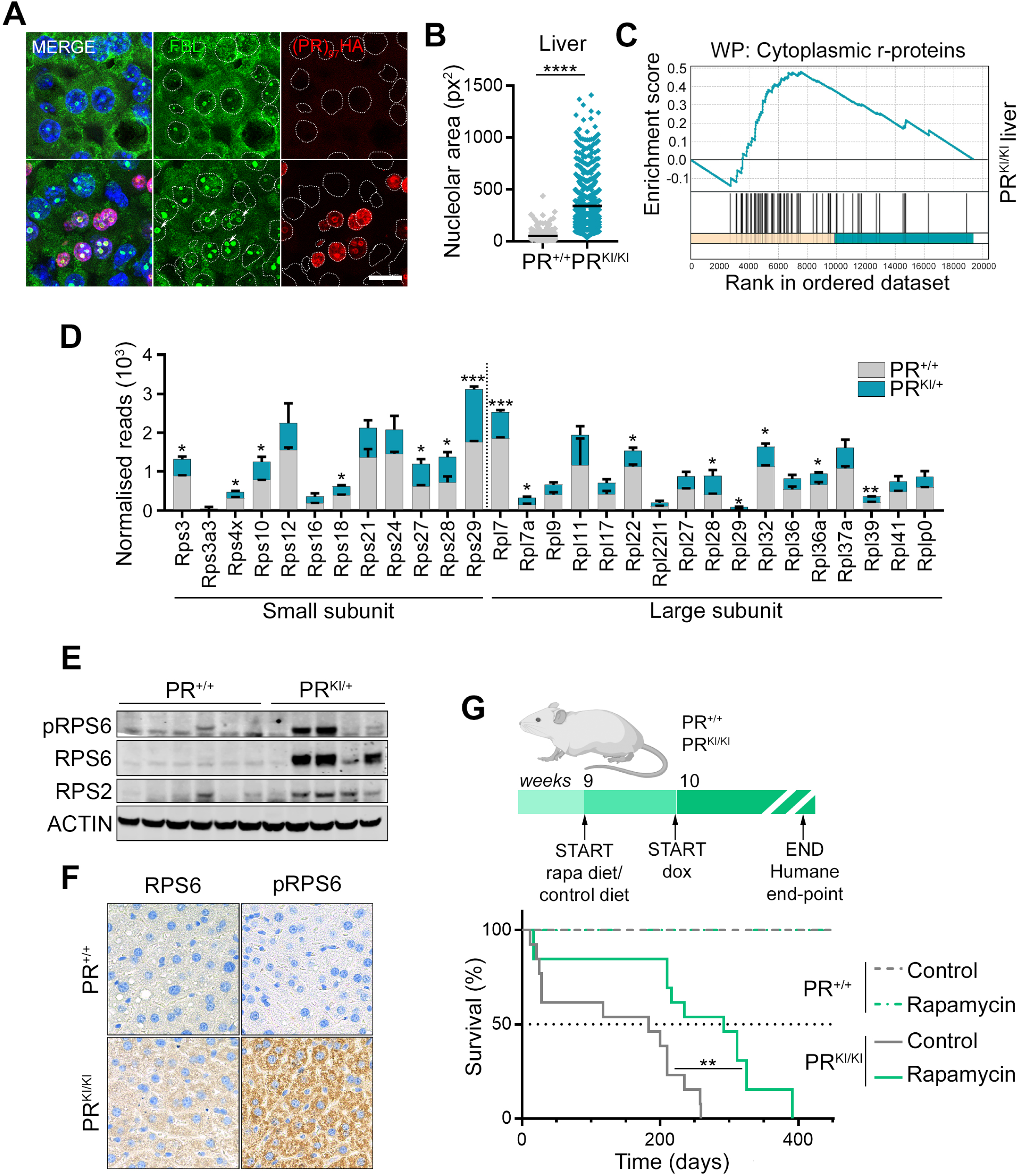
NS and mTOR hyperactivation contribute to the progeria of (PR)_97_^HA^ expressing mice. (A) IF of (PR)_97_^HA^ (red) and FBL (green), in liver sections from littermate PR^+/+^ and PR^KI/KI^ mice, 3 weeks after treatment with dox in the drinking water. Scale bar (white) represents 10 μm. (B) Quantification of nucleolar FBL+ area from the images shown in (A). (C) GSEA enrichment plots for the “cytoplasmic r-proteins” class from RNA-seq data comparing liver RNA from PR^+/+^ and PR^KI/+^ littermates after 17 weeks of treatment with dox. (D) Normalised read counts of all detected r-protein mRNAs, quantified by RNA-seq in total RNA of livers from non-inducible (grey, PR^+/+^) and inducible (blue, PR^KI/+^) littermates, after 17 weeks of treatment with dox. (E) WB evaluating the levels of mTOR signalling (pRPS6), and of total RPS6 and RPS2 levels in livers from non-inducible PR^+/+^ and inducible PR^KI/+^ littermates after 20 weeks of treatment with dox. (F) RPS6 and pRPS6 IHC in liver sections from a non-inducible mouse (PR^+/+^) and an inducible littermate (PR^KI/KI^), after 3 weeks of treatment with dox. (F) Diagram illustrating the experimental pipeline for the treatment of PR^+/+^ and PR^KI/KI^ animals with rapamycin and dox. (G) Kaplan-Meyer representation of the lifespan of non-inducible PR^+/+^ (*n* = 13), and inducible PR^KI/KI^ (*n* = 13) littermates. Animals started treatment with rapamycin in their diet at week 9, and the treatment with dox started on the following week. Note that rapamycin extends the median lifespan by 60% in inducible mice. (** *P* < 0.01; Mantel-Cox test). Data information: *P* > 0.05; * *P* < 0.05; ** *P* < 0.01; *** *P* < 0.005; \*\*\*\**P* < 0.0001; *t*-test.

In addition to the increase in total r-protein levels, WB and IHC revealed higher levels of RPS6 phosphorylation, consistent with hyperactivation of the mTOR pathway in (PR)_97_^HA^ -expressing livers (**Fig. 5E, F**). Finally, Rapa treatment increased the viability of dox-treated PR^KI/KI^ MEFs (**Fig. S8C-E**) but also, importantly, prolonged survival of dox-treated PR^KI/KI^ mice (**Fig. 5G**). Noteworthy, a similar increase in the mTOR pathway and r-proteins was also observed by analysing transcriptomic data of human motor neurons derived from induced pluripotent stem cells of *C9ORF72* ALS patients (**Fig. S9**), available at the NeuroLINCS repository (http://neurolincs.org)^38^. Together, these experiments demonstrate that (PR)_n_ expression *in vivo* leads to NS, mTOR hyperactivation and an accumulation of free r-proteins, which induces accelerated ageing and premature death, that can be alleviated by mTOR inhibition.

### The accumulation of free r-proteins is a common outcome of NS

Based on our observations with (PR)_n_ peptides, we wondered if the accumulation of free r-proteins was a common hallmark of cells suffering from NS. To test this hypothesis, we repeated our proteomic experiments from sucrose gradients and analysed the fraction of the proteome that was not bound to ribosomes in cells experiencing NS triggered by different insults. The sources of NS were: (1) treatment with Act D; (2) depletion of the RNA Pol I associated factor TIF-IA, previously shown to induce NS *in vitro* and *in vivo,* and (3) inducible expression of a HGMB1 variant harbouring an extra C-terminal arginine-rich peptide, recently reported to abnormally accumulate at nucleoli and identified as a driver mutation in a yet-to-be-named rare human disorder^39^. Strikingly, all these proteomic analyses showed a significant enrichment of ribosome-free r-proteins in all cases, which could also be detected by IF (**Fig. 6**). Hence, the accumulation of free r-proteins is a hallmark of cells suffering from NS.

**Figure 6.**
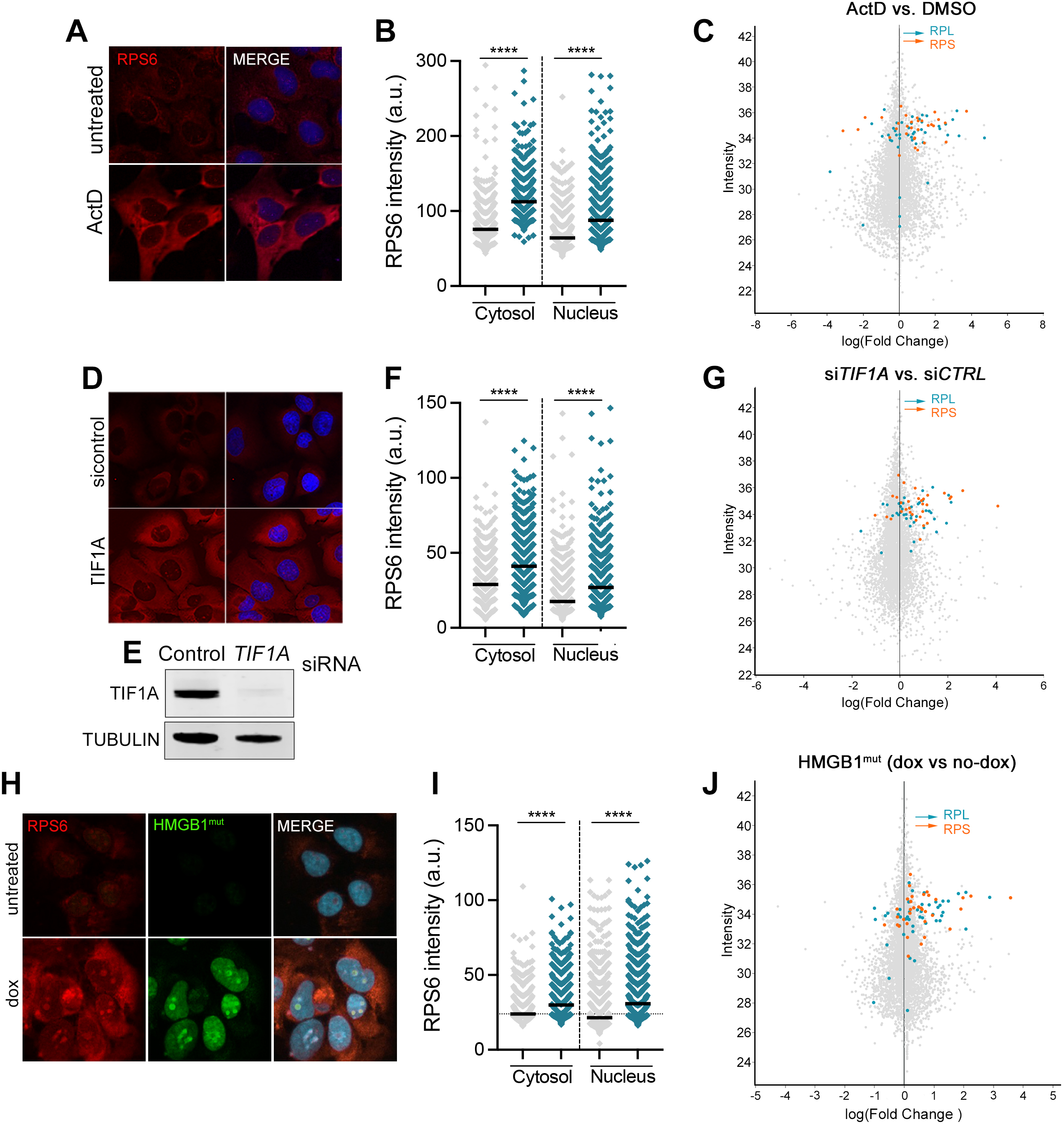
The accumulation of free r-proteins is a common outcome in response to NS. (A) RPS6 (red) IF, in U2OS cells treated with ActD (5nM) for 24 hrs. (B) HTM-dependent quantification of RPS6 levels from data shown in (A). (C) Proteomic analysis of the polysome-free fraction from U2OS cells treated as in (A). Note the enrichment of free RPL (blue) and RPS (red) r-proteins in Act D-treated cells. (D) RPS6 (red) IF, in U2OS cells transfected with a *TIF1A*-targeting siRNA (or control siRNA) for 24 hrs. (E) WB illustrating the efficient TIF1A depletion. TUBULIN was used as a loading control. (F) HTM-dependent quantification of RPS6 levels from data shown in (D). (G) Proteomic analysis of the polysome-free fraction from U2OS cells treated as in (D). Note the enrichment of free RPL (blue) and RPS (red) r-proteins in TIF1A-depleted cells. (H) RPS6 (red) IF, in U2OS cells upon dox-dependent expression of a mutant form of HMGB1 harbouring a C-terminal R-rich insertion (HMGB1^mut^, green) for 48 hrs. As previously reported ^39^, HMGB1^mut^ accumulated at nucleoli which increased their size. (I) HTM-dependent quantification of RPS6 levels from data shown in (H). (J) Proteomic analysis of the polysome-free fraction from U2OS cells treated as in (H). Once again, note the enrichment of free RPL (blue) and RPS (red) r-proteins in U2OS cells upon expression of HMGB1^mut^.

## Discussion

Ribosome biogenesis is the most energy-demanding activity in a cell^40^. As a consequence, perturbing nucleolar function can have a profound impact on cell fitness and/or viability. Correspondingly, abnormalities in nucleolar activity or structure, now collectively known as NS, have often been found in tissues from patients of several human diseases such as ribosomopathies, cancer or neurodegeneration^17^. Very recent work added brachyphalangy, polydactyly and tibial aplasia/hypoplasia syndrome (BPTAS), to the type of diseases associated to nucleolar dysfunction^39^. Interestingly, mutations in BPTAS patients lead to the generation of C-terminal arginine-rich insertions in several proteins, all of which accumulate at nucleoli, and alter their function and phase separation properties. These mutations are reminiscent of those found in patients of neuromuscular pathologies with mutations in *C9ORF72* or *SCA36*, which also lead to the generation of arginine-rich peptides that accumulate at nucleoli and perturb ribosome biogenesis^23^. However, a mechanistic understanding of how these mutations lead to pathology and of strategies that can potentially alleviate it, is missing.

Initial evidences revealed that one way by which NS can kill cells is by activating the stress-sensor P53^15^. Subsequent work indicated that P53 activation by NS is triggered by specific ribosomal proteins such as RPL5 or RPL11, which interact with P53 ubiquitin ligases MDM2 and MDM4^8^. However, this model is necessarily incomplete as organisms like *Drosophila melanogaster* lack MDM2 orthologues and are still significantly affected by NS^41^. Other pathways like the Jun N-terminal kinase (JNK)^42^, the retinoblastoma tumor suppressor (Rb)^43^ or NF-kB^44^ have been also been suggested to contribute to the toxicity of NS. Besides the activation of a specific pro-apoptotic signaling cascade, an alternative model is that NS triggers a systemic metabolic imbalance that compromises cell viability. Along these lines, a very interesting recent study in flies revealed that, contrary to the original hypotheses, the pathologies found in ribosomopathies are not due to defects in mRNA translation, but rather to the widespread accumulation of free r-proteins, which due to their abundance end up clogging clearance mechanisms such as the proteasome or autophagy^45^.

Our work extends on these findings, and reveals that the accumulation or ribosome-free r-proteins is not restricted to ribosomopathies but is rather a common outcome that occurs in response to NS, regardless of the insult. Moreover, and as already suggested for ribosomopathies, our data suggest that the use of mTOR inhibitors as a general strategy to counteract the toxicity of NS. We are nevertheless aware of the limitations of inhibiting mTOR function in humans, as its effects are not restricted to the activation of autophagy but also impair other important aspects such as insulin or growth factor signaling. Thus, there is a clear need to identify more selective ways of stimulating the clearance of free r-proteins, and identifying specific inductors of ribophagy emerges as an interesting alternative. Finally, and besides increasing our understanding on how NS kills cells, our research adds support for the emerging links between nucleoli and ageing^30,46^, being to our knowledge the first example to document NS-triggered accelerated aging in mammals.

## Materials and Methods

### Cell Culture

All cells were cultivated at 37°C, in a humidified air atmosphere with 5% CO_2_, unless otherwise specified. U2OS (ATCC), NSC-34 (ATCC), U2OS-TRex(PR)^97^, inducible U2OS-HMGB1mut, and NSC-34 TRex(PR)^97^cells were cultivated in standard high glucose Dulbecco’s Modified Eagle Medium (DMEM) supplemented with 10% fetal bovine serum (FBS), 2 mM L-glutamine and 1% penicillin/streptomycin, using tetracycline-free FBS (PAN biotech, P30-3602) in the case of inducible cells. Mouse embryonic fibroblasts (MEFs) were cultivated in DMEM supplemented with 15% FBS, and under hypoxic conditions (5% CO_2_, and 5% O_2_). Immortalized *Tsc^+/+^* and *Tsc2^-/-^* MEF were a kind gift from Alejo Efeyan (CNIO, Madrid). MEFs from PR^KI/KI^ mice were generated from mouse embryos extracted at day 14 post-coitum.

The dox-inducible NSC34(PR)^97^ cell line was generated by transfecting NSC34-TRex cells with a pINTO-C-(PR)_97_-HF plasmid, as previously described^47^. Dox-inducible GFP-HMGB1mut U2OS cells were generated using a previously described pPB-TetON-mEGFP-HMGB1-MUT PiggyBac transposon (Addgene #194562)^39^, and a transposase expressing pCMV-HyPBase plasmid (kind gift from Alberto Pendás, CIC Salamanca). Spontaneously resistant NSC-34^R^ clones were generated from wild-type NSC34 cells. 4,000 cells were seeded per well of a 6 well plate. On the next day, cells were treated with 5 μM of recombinant (PR)_20_ peptides (with a C-terminal HA epitope tag, Genscript) for 10 days, changing medium supplemented with fresh (PR)_20_ every 2 days. At this time, (PR)_20_ resistant cells were trypsinized and dispensed as single cells in 96 well plates by FACSAria^TM^ III (BD Biosciences). The cells were subsequently exposed to 10 µM (PR)_20_ for 2 – 3 weeks, with the medium being changed each week. Resistant clones were selected for further analuses

For NSC34 differentiation, cells were seeded onto plates containing DMEM supplemented with 10% FBS, 2 mM L-glutamine and 1% penicillin/streptomycin. On the next day, media was changed to Neurobasal medium (Thermo Fisher scientific, 2121103049) containing B-27 supplement (Thermo Fisher scientific, 17504044) for 48 h.

### Cell viability

For U2OS(PR)^97^ cells, viability was quantified by high throughput microscopy (HTM). 5,000 cells were seeded, and treated with 1 µL/ mL dox, alone or with other treatments. After the indicated time, cells were fixed with 4% paraformaldehyde (PFA) and stained with DAPI. Images were automatically acquired from each well using an Opera High-Content Screening System (Perkin Elmer). A 10x magnification lens was used and images were taken at non-saturating conditions. Images were segmented using DAPI signals to count the number of cells. For NSC34(PR)^97^ cells, viability was quantified by manual counting of live cells. 100,000 cells were seeded on gelatin pre-treated 12-well plates, and on the following day treated with 1 µg/mL dox with 50 nM rapamycin, or 50 nM torin-1. 72 hours later cells were trypsinized and counted, using trypan blue to identify dead cells. To test the response to different drugs, 10,000 NSC34^WT^ and NSC34^R^ were seeded, and treated with the following compounds AZ20 (ATRi), 20 µM, cisplatin, 25 µM, camptothecin (CPT), 8 µM, (PR)_20_, 20 µM, Actinomycin D (ActD), 5 nM, and CX-541, 10 µM on the following day. Viability was quantified 24 hours after treatment using a luminescent system (CellTiter-Glo, Promega). In experiments using MEFs, 2000 cells were seeded, and treated with dox, with or without rapamycin (1 nM), on the following day. Viability was quantified 48 hours after treatment using a luminescent system (CellTiter-Glo, Promega). For clonogenic assays, U2OS(PR)^97^ cells were simultaneously treated with 30 nM rapamycin or 3 nM torin-1, with dox, for 48 h, and then allowed to grow for 10 days before fixing. At the end of the experiments, cells were fixed and stained with 0.4% methylene blue in methanol for 30 min.

### Immunofluorescence and high throughput microscopy

For immunofluorescence, cells were fixed with 4% PFA prepared in PHEM buffer (60 mM Pipes, 25 mM Hepes, 10 mM EGTA, 2 mM MgCl_2_ pH 6.9) containing 0.2% of Triton X-100, and permeabilized with 0.5% Triton X-100 after fixation. For high throughput microscopy, cells were grown on µCLEAR bottom 96-well plates (Greiner Bio-One) and immunofluorescence of RPS6 (Cell Signaling, 2217), HA-tag (Roche, 11867423001), p53 (Santa Cruz sc-126), TUBB3 (Biolegend, 801202), FBL (Cell Signaling, 2639), UBF (Santa Cruz, sc-13125) was performed using standard procedures. Analysis of transcription by EU and translation by HPG incorporation were done using Click-It kits (Invitrogen) following the manufacturer’s instructions. In all cases, images were automatically acquired from each well using an Opera High-Content Screening System (Perkin Elmer). A 20x or 40x magnification lens was used and images were taken at non-saturating conditions. Images were segmented using DAPI signals to generate masks matching cell nuclei, from which the mean signals for the rest of the stainings were calculated.

### Polysome analyses

Exponentially growing U2OS cells were transfected with 50 nM of control siRNAs or siRNAs targeting human *TIF1A* (Horizon Discovery biosciences, ON-TARGETplus siRNAs), or treated with 5 nM Act D (Sigma). Inducible U2OS(PR)^97^ and U2OS-HMGB1mut cells were treated with dox (Sigma) for 48 h. Ribosomes were stalled by addition of 100 µg/mL cycloheximide (CHX) for 5 min, and cells lysed in polysome lysis buffer (15 mM Tris-HCl pH 7.4, 15 mM MgCl_2_, 300 mM NaCl, 1% Triton-X-100, 0.1% β-mercaptoethanol, 200 U/mL RNAsin (Promega), 1 complete Mini Protease Inhibitor Tablet (Roche) per 10 mL). Nuclei were removed by centrifugation (9300×G, 4°C, 10 min) and the cytoplasmic lysate was loaded onto a sucrose density gradient (17.5–50% in 15 mM Tris-HCl pH 7.4, 15 mM MgCl_2_, 300 mM NaCl and, for fractionation from BMDM, 200 U/mL Recombinant RNAsin Ribonuclease Inhibitor, Promega). After ultracentrifugation (2.5 h, 35,000 rpm at 4°C in a SW60Ti rotor), gradients were eluted with a Teledyne Isco Foxy Jr. system into 13 fractions of similar volume. The first two, ribosome-free fractions, were compared with the rest of polysome fractions.

For proteomic analyses, samples from the sucrose gradient fractions were solubilized in 2% SDS, 100 mM TEAB pH 7.55. Proteins were reduced and alkylated (15 mM TCEP, 25 mM CAA) 1h at 45 °C in the dark. Then, samples were digested following the solid-phase-enhanced sample-preparation (SP3) protocol^48^. Briefly, ethanol was added to the samples to a final concentration of 70% and proteins were incubated for 15 minutes with SP3 beads at a bead/protein ratio of 10:1 (wt/wt). Beads were rinsed using 80% EtOH and proteins were digested with 100 µL of trypsin in 50 mM TEAB pH 7.55 (Promega, protein:enzyme ratio 1:50, 16 h at 37 °C). Resulting peptides were desalted using C18 stage-tips. LC-MS/MS was done by coupling an UltiMate 3000 RSLCnano LC system to a Q Exactive HF mass spectrometer (Thermo Fisher Scientific) or to a Q Exactive Plus mass spectrometer (Thermo Fisher Scientific). Peptides were loaded into a trap column (Acclaim™ PepMap™ 100 C18 LC Columns 5 µm, 20 mm length) for 3 min, at a flow rate of 10 µL/min in 0.1% FA. Then, peptides were transferred to an EASY-Spray PepMap RSLC C18 column (Thermo Fisher Scientific) (2 µm, 75 µm x 50 cm) operated at 45 °C and separated using a 60 min effective gradient (buffer A: 0.1% FA; buffer B: 100% ACN, 0.1% FA) at a flow rate of 250 nL/min. The gradient used was, from 2% to 6% of buffer B in 2 min, from 6% to 33% B in 58 minutes, from 33% to 45% in 2 minutes, plus 10 additional minutes at 98% B. Peptides were sprayed at 1.5 kV into the mass spectrometer via the EASY-Spray source and the capillary temperature was set to 300 °C.

The Q Exactive HF mass spectrometer was operated in a data-dependent mode, with an automatic switch between MS and MS/MS scans using a top 15 method. (Intensity threshold ≥ 6.7E4, dynamic exclusion of 25 s and excluding charges +1 and > +6). MS spectra were acquired from 350 to 1400 m/z with a resolution of 60,000 FWHM (200 m/z). Ion peptides were isolated using a 2.0 Th window and fragmented using higher-energy collisional dissociation (HCD) with a normalised collision energy of 27. MS/MS spectra resolution was set to 15,000 (200 m/z). The ion target values were 3e6 for MS (maximum IT of 25 ms) and 1E5 for MS/MS (maximum IT of 15 ms). The Q Exactive Plus mass spectrometer was also operated in a data-dependent mode, with an automatic switch between MS and MS/MS scans using a top 15 method. (Intensity threshold ≥ 56.7E4, dynamic exclusion of 25 s and excluding charges +1 and > +6). MS spectra were acquired from 350 to 1400 m/z with a resolution of 70,000 FWHM (200 m/z). Ion peptides were isolated using a 2.0 Th window and fragmented using higher-energy collisional dissociation (HCD) with a normalised collision energy of 27. MS/MS spectra resolution was set to 17,500 (200 m/z). The ion target values were 3e6 for MS (maximum IT of 25 ms) and 1E5 for MS/MS (maximum IT of 45 ms).

Raw files were processed with MaxQuant (v 1.6.0.43) using the standard settings against a human protein database (UniProtKB/Swiss-Prot, 20,373 sequences). Carbamidomethylation of cysteines was set as a fixed modification whereas oxidation of methionines and protein N-term acetylation were set as variable modifications. Minimal peptide length was set to 7 amino acids and a maximum of two tryptic missed-cleavages were allowed. Results were filtered at 0.01 FDR (peptide and protein level). Afterwards, the “proteinGroups.txt” file was loaded in Prostar^49^ using the intensity values for further statistical analysis. Briefly, proteins with less than two valid values in at least one experimental condition were filtered out. Missing values were imputed using the SLSA algorithms^50^ for partially observed values and DetQuantile for values missing on an entire condition. Differential analysis was done using the empirical Bayes statistics Limma. Proteins with a p.value < 0.05 and a log2 ratio >1 or <-1 were defined as regulated. The FDR was estimated to be below 7.43% by Benjamini-Hochberg.

### Generation of PR^KI^ mice

The inducible PR^KI^ mouse line was generated by using a previously system^37^ that enables site-specific recombination targeted to the *Col1A1* locus. The (PR)_97_-HA-FLAG cassette was extracted from pINTO-C-(PR)_97_-HF by PCR amplification, and cloned into the flp-in pBS31, using the EcoRI and MluI sites. The circular pBS31-(PR)_97_-HF plasmid was co-electroporated into KH2 mESCs (hybrid B6.129 background), which harbour the rtTA transactivator in the widely expressed *Rosa26* locus, along with a vector expressing the FLPe recombinase. After selection in hygromicin 200 µg/mL, single mES clones were isolated, and functionally tested with dox treatment *in vitro*. A clone showing high and homogeneous induction levels of (PR)_97_ was selected, microinjected into E2.5 C57BL/6_TyrC morulae, and transferred to CD1 foster females, giving rise to chimeric animals. A male with a high degree of chimaerism was selected and backcrossed with C57BL/6 females to test for germ line transmission. The resulting inducible mice in this study are in a mixed C57BL6.129 background.

DNA was extracted from ear tissue samples using isopropanol precipitation, following standard protocols, and used in PCRs to determine the genotypes of the mice. The primer sequences are as follows (5’-3’): Rosa26^F^ AAAGTCGCTCTGAGTTGTTAT, Rosa26^R-WT^ GGAGCGGGAGAAATGGATATG, Rosa26^R-KI^ GCGAAGAGTTTGTCCTCAACC, Col1A1^F^ CCCTCCATGTGTGACCAAGG, Col1A1^R-WT^ GCACAGCATTGCGGACATGC and Col1A1^R-KI^ GCAGAAGCGCGGCCGTCTGG. The Rosa26 locus yields a wild-type band of 500bp and 300bp for the knock-in allele, while the Col1A1 wild-type allele gives a band of 331 bp, and the knock-in of 551 bp.

For the initial characterization, mice were treated, starting at 5 weeks, with oral dox (PanReac), 2 mg/mL, in 5% sucrose (Sigma), to make the solution palatable. Teklad Global 18% Protein Rodent Diet chow diet (2018S, Harlan Laboratories) was freely available. Encapsulated rapamycin was administered in the diet. Rapamycin encapsulated in eudragit, or empty eudragit microcapsules (Rapamycin holdings), were mixed with a standard 5LG6 ModLabDiet (TestDiet) at 425 ppm, and irradiated, to make the rapamycin diet, and the control diet. Standard chow was switched to rapamycin, or control diet, at 9-10 weeks of age, and the mice were allowed to adapt for another week before beginning the dox treatment, in which sucrose was replaced by 0.4% saccharin (Sigma), to reduce free sugar intake. The treatments were postponed until this age to allow for weight-loss caused by the rapamycin diet. Sucrose was eliminated to prevent rapamycin-induced diabetes. Health status of mice was monitored daily, and weight every 1-2 weeks. Mice were maintained at the CNIO animal facility under standard housing conditions with free access to chow diet and water, following the guidelines of the Federation of European Laboratory Animal Science Association. All mouse work was performed in accordance to the Guidelines for Humane Endpoints for Animals Used in Biomedical Research and under the supervision of the Ethics Committee for Animal Research of the “Instituto de Salud Carlos III”.

### In vivo imaging

For microCT scans mice were anaesthetized with a continuous flow of 1% to 3% sevoflurane/oxygen mixture (2 L/min), and the whole body was able to be imaged at one time using the eXplore Vista micro-CT scanner (GE Healthcare, London, Canada). The isotropic resolution of this instrument is 45 µm. The micro-CT image acquisition consisted of 400 projections collected in one full rotation of the gantry in approximately 10 min. The X-ray tube settings were 80 kV and 450 µA. The angles were measured using the software 3D slicer. The kyphosis angle was determined between the C1-T3-T9 vertebrae in each animal.

### Western Blotting

Cell pellets were recovered, and washed with cold PBS, before lysis at 4°C, on a shaker, using Urea buffer (50 mM Tris, pH 7.5, 8 M urea, and 1% CHAPS). To obtain mouse tissue protein extracts, a sample of the tissue was homogenized using zirconium beads in RIPA buffer (50 mM Tris, pH 8.0, 0.2% nonidet-P40, 200 mM NaCl, 50 µM glicerolphosphate, and 1% tween-20) with protease and phosphatase inhibitors. The protein fraction was isolated by centrifugation at 13.2 rpm, for 15 minutes, and quantified using the Bio-Rad Protein Assay (Bio-Rad). Approximately 20 µg of sample was mixed with NuPAGE LDS (LifeTechnologies) and 10 mM dithiothreitol (DTT) (Sigma), and incubated at 70°C for 10 minutes. The extracts were resolved in precast 4-20% gradient polyacrylamide gels (Invitrogen), and transferred using standard methods. After blocking, the membrane was incubated overnight at 4°C with the primary antibody, and 1 h at room temperature with the secondary. A complete list of all antibodies used can be found in Table 5. Fluorophore-conjugated secondary antibodies were used for detection, using the Li-Cor LCx system. The following primary antibodies were used: RPS6 (Cell Signaling 2217), p-RPS6 (Cell Signaling, 2215), HA-tag (Roche, 11867423001), p53 (Santa Cruz sc-126), actin (Sigma, A5441), S6K1 (Cell Signalling, 2708), p-S6K1 (Cell Signalling, 2215), TIF1A (Santa Cruz, sc-271266) and RPS2 (Santa Cruz, sc-130399).

### RNA-seq

Total RNA was extracted from liver tissue using TRIzol reagent (Invitrogen), according to the manufacturer’s protocols. A portion of the RNA was purified a second time with the RNA Clean & Concentrator-5 kit (Zymo Research), according to manufacturer’s instructions, and submitted to standard quality controls. The sequencing library was constructed with the QuantSeq 3’ mRNA-Seq Library Prep Kit (Lexogen), and approximately 10 million reads were obtained by Illumina sequencing. Differential expression analysis was performed using Bluebee® (Lexogen). Functional analysis was done by Gene Set Enrichment Analysis (GSEA) of the expression data, according to the publishers’ instructions^51^.

### qRT-PCR

RNA was extracted from NSC34 cells using the Absolutely RNA Microprep kit (Agilent). The rRNA levels were measured by real-time quantitative PCR after reverse transcription of RNA, using the SuperScript III Platinum SYBR Green One-Step qRT-PCR Kit with ROX (Invitrogen), and 3 ng of total RNA. Each value was normalized against that of *Eif1* of the corresponding line. The sequences of the primers used are as follows: *45S* F GGCTGGGGTTGGAAAGTTTC, *45S* R CAAGGGCATTCTGAGCATCC, *18S* F CTGGATACCGCAGCTAGGAA, *18S* R GAATTTCACCTCTAGCGGCG, *5.8S* F GTCGATGAAGAACGCAGCTA, *5.8S* R AACCGACGCTCAGACAGG, *28S* F CGGCGGGAGTAACTATGACT, *28S* R GCTGTGGTTTCGCTGGATAG, *Eif1* F TGGTACTGTAATTGAGCATCCAG, *Eif1* R CCTTAGCCAGCCCAATCTCT. All reactions were carried out in MicroAmp© Optical 384-Well plates (Applied Biosystems) in the QuantStudio™ 6 Flex Real-Time PCR System (Thermo Fisher) using standard protocols.

### Immunohistochemistry and immunofluorescence

Tissues were fixed in formalin and embedded in paraffin for subsequent processing. Sections of 2.5 µm were treated with citrate for antigen retrieval, and processed for immunohistochemistry with hematoxylin and eosin staining, or HA-tag antibodies, following standard protocols. Images were captured with a Leica DM2000 LED optical microscope, using a 20x lens. Immunofluorescence on tissue sections was carried out as previously described^52^. Briefly, after antigen retrieval, the sections are rinsed, and permeabilized with 0.25% Triton X100 and 0.2% gelatin in PBS, after which they are blocked in 5% BSA in permeabilization buffer. They were incubated with the primary antibodies (anti-HA, Roche, 11867423001; anti-FBL, Cell Signaling, 2639; anti-PR, a kind gift from Giovanna Roncador) overnight, at 4°C, and secondary for 1 h at room temperature. Finally, after rinsing, the sections are incubated in 10 mM CuSO_4_/50 mM NH_4_Cl solution, dried, and mounted with Fluoromount-G (SouthernBiotech) mounting media. Images were captured with a Leica SP5 WLL confocal microscope. A 40x magnification lens was used and images were taken at non-saturating conditions. Images were segmented using DAPI and 488-FBL staining signals to generate masks matching cell nuclei, and nucleoli, respectively, to obtain the nucleolar area.

### Drug set enrichment analysis

The CMap Query clue.io tool (https://clue.io/)^36^ was used to extract perturbagens (compounds, over-expression, knock-out or knock-down of genes) with signatures similar to spontaneously resistant NSC34^R^ cells. For this, 50-100 genes upregulated in the proteomics profile were used as an input for the Query CMap tool, employing Gene expression (L1000), Touchstone and individual query as query parameters. Similarity scores were downloaded and results were sorted based on their scores and the type of perturbagen.

DSEA was carried out, as previously described^53^. Briefly, the GSEA method implemented in the R package fgsea^54^ was adapted to enable enrichment analyses of “drug classes” based on their mechanism of action. The CMap compound similarity scores were ranked, and the GSEA method was applied to the ranked list with drug sets for DSEA analysis. The drug classes were based on the mechanism of action, which was available in the annotation data included in CMap analysis in the “description” field. For the computational analysis, R version 3.6.3 was used. A sample of the code used for DSEA analyses can be obtained from https://github.com/Genomic-Instability-Lab/An-in-silicoanalysis-of-drugs-potentially-modulating-the-cytokine-storm-triggered-by-SARS-CoV-2-inf.

### NeuroLINCS data analysis

Expression data from motor neurons, differentiated from induced pluripotent stem cells of ALS patients and unaffected subjects, was obtained from the NeuroLINCS data repository (Leslie Thompson: diMN (Exp 3) - ALS and Control (unaffected) diMN cell lines differentiated from iPS cell lines using a short and direct differentiation protocol - RNA-seq, 2017, LINCS (dataset), http://identifiers.org/lincs.data/LDS-1499, ; retrieved: 21/07/2021). Differential expression analysis between unaffected subjects and sporadic ALS cases, or unaffected subjects and C9ORF72-ALS cases, was carried out using the limma-voom tool^55^, available on the Galaxy platform (https://usegalaxy.org/). The P-value of enrichment of mTORC1 signaling and of r-proteins was obtained by a hypergeometric test.

### Data Availability

Mass spectrometry data have been deposited to the ProteomeXchange Consortium via the PRIDE partner repository with the dataset identifier PXD043752. The following username and password (pqPTpLFK) can be used during the peer review purposes. RNA-seq data are available at the GEO repository with accession number GSE239319.

#### Statistics

All statistical analyses were performed using Prism software (GraphPad Software) and statistical significance was determined where the p-value was <0.05 (*), <0.01 (**), <0.001 (***) and <0.0001 (****). Survival data was evaluated using Kaplan-Meier analyses.

## Acknowledgments

The authors want to help the proteomics, genomics and transgenics units of the CNIO for their technical support in this study. O.F-C. is supported by grants from the Spanish Ministry of Science, Innovation and Universities (PID2021-128722OB-I00, co-financed with European FEDER funds), the Spanish Association Against Cancer (AECC; PROYE20101FERN) and La Caixa Foundation (HR22-00890). The authors declare no competing financial interests.

## Author Contributions

O.S. contributed to most of the experiments; B.J. generated NSC34^R^ cell lines; I.V. helped with polysome fractionation experiments; V.L. contributed to experiments, helped to supervise the work and worked on the manuscript; O.F. conceived the experimental plan, coordinated the study, supervised the experiments and wrote the manuscript.

