## Supplementary material for "Nucleolar stress caused by arginine-rich peptides triggers a ribosomopathy and accelerates ageing in mice": Figures S1-S9

**Figure S1. Death by (PR)n peptides is not additive with other sources of NS.** (A, B) Viability of U2OS$^{(PR)97}$ cells, as measured by HTM-mediated quantification of nuclei stained with DAPI, after 48 hrs of treatment with the NS inducers CX-5461 (A) or ActD (B) in the presence (green) or absence (grey) of dox. (C, D) Viability of two biologically independent conditional heterozygous *Rpl11*$^{+/lox}$ MEFs, as measured by HTM-mediated quantification of nuclei stained by DAPI, after 48 hrs of treatment with $(PR)_{20}$ peptides. Deletion of the conditional (lox) *Rpl11* allele was done by addition of 4-hydroxy-tamoxifen (OHT), as all cells expressed an OHT-inducible Cre$^{ER}$ fusion protein.

*Sirozh et al* **FigS1**

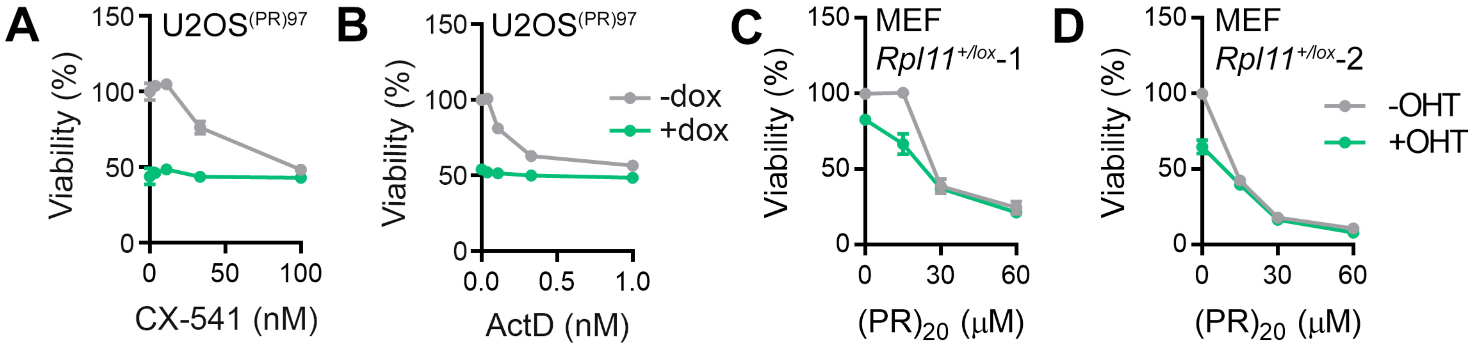

**Figure S2. Persistent exposure to (PR)n peptides triggers P53-independent toxicity.** (A) P53 WB in U2OS$^{(PR)97}$ cells treated with dox for 0, 24, 48 and 72 hrs, or ActD for 24 hrs. In contrast to ActD, $(PR)_{97}$ expression triggers a limited activation of P53. (B) Representative immunofluorescence of P53 (green) in U2OS$^{(PR)97}$ cells 48 hrs after treatment with dox. Scale bar (white) represents 10 $\mu$m. (C) HTM-dependent quantification of the P53 signal from (B). Black lines indicate mean values. ****$P < 0.0001$; *t*-test. (D) Clonogenic survival assay of wild-type and P53-null HCT-116 cells, exposed to $(PR)_{20}$. HCT-116 cells were treated with 15 $\mu$M $(PR)_{20}$, for 72 hrs, and then allowed to grow for 7 days before fixing. (E) WB confirming the depletion of P53 in U2OS$^{(PR)97}$. P53 depletion was performed using CRISPR after transfecting Cas9-expressing U2OS$^{(PR)97}$ cells with control or *P53*-targeting sgRNAs. (F) High density survival assay of wild-type and the pool of P53-null U2OS$^{(PR)97}$ cells, exposed to doxycycline for 72 hrs.

*Sirozh et al* **FigS2**

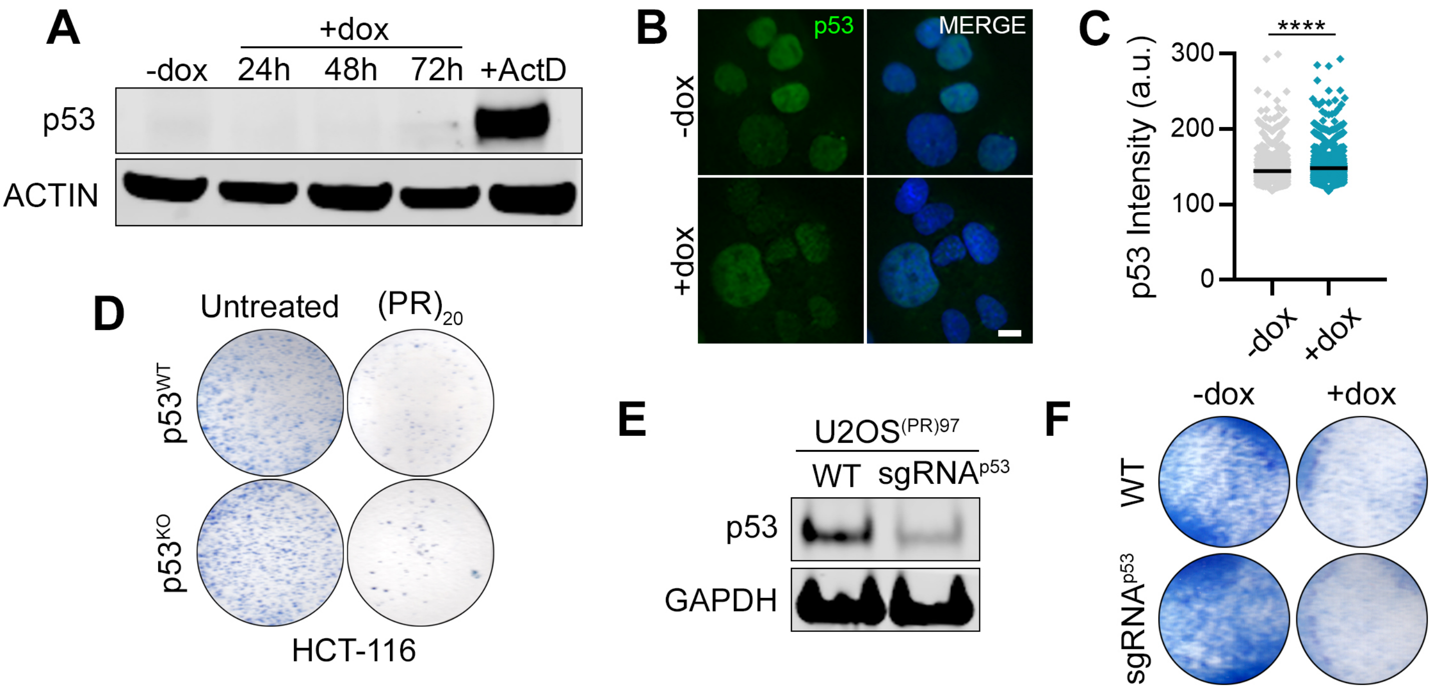

**Figure S3. Polysome RPL depletion and overall accumulation of r-proteins upon (PR)$^{97}$ expression.** Protein extracts from U2OS$^{(PR)97}$ cells, after a 48 hrs of treatment with dox were fractionated in sucrose gradients and compared to fractions from untreated cells. The fold change in the polysome fraction is represented in the *y* axis, while the fold change in the ribosome-free fraction is represented in the *x* axis. r-proteins are marked with orange (RPS) and blue (RPL).

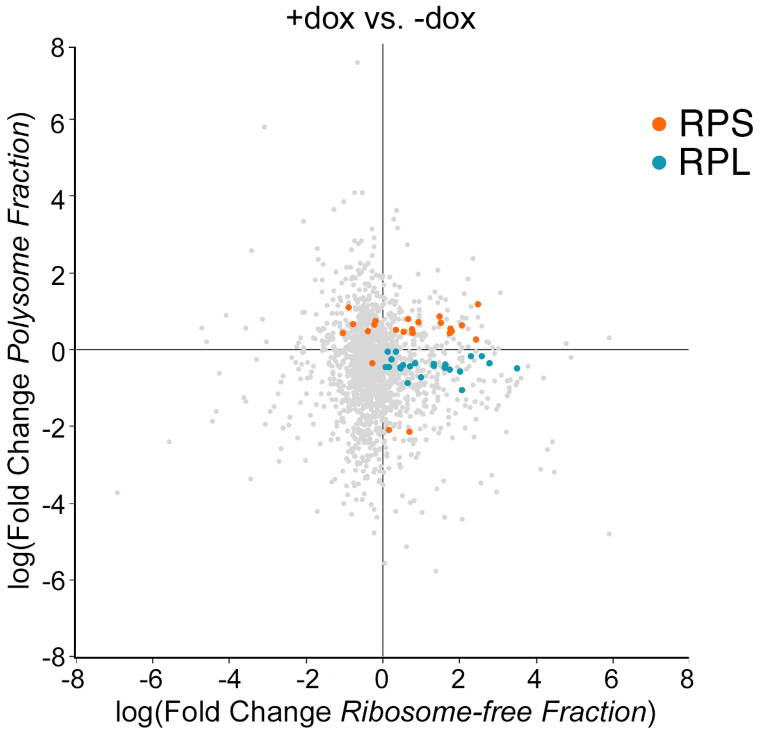

+dox vs. -dox
RPS
RPL
log(Fold Change *Polysome Fraction*)
log(Fold Change *Ribosome-free Fraction*)
8
6
4
2
0
-2
-4
-6
-8

**Figure S4. Reduced translation and ribosome biogenesis in (PR)n-resistant cells.** (A) Representation of statistically significant GO terms of downregulated biological processes involved in translation and ribosome biogenesis, as detected in GSEA analyses from the proteomic comparison of NSC34$^{R2}$ with wild-type NSC34$^{WT}$ cells. The normalised enrichment score is indicated by the bars, while the p-value is colour-coded according to the legend. (B) Volcano plot of all proteins identified by the proteomic comparison of NSC34$^{WT}$ with NSC34$^{R2}$ cells. r-proteins are marked with orange (40S) and blue (60S). (C, D) GSEA enrichment plots for the gene ontology classes corresponding to ribosome biogenesis (C), and translation initiation (D) derived from RNA-Seq data comparing NSC34$^{R2}$ cells with the parental NSC34 cell line.

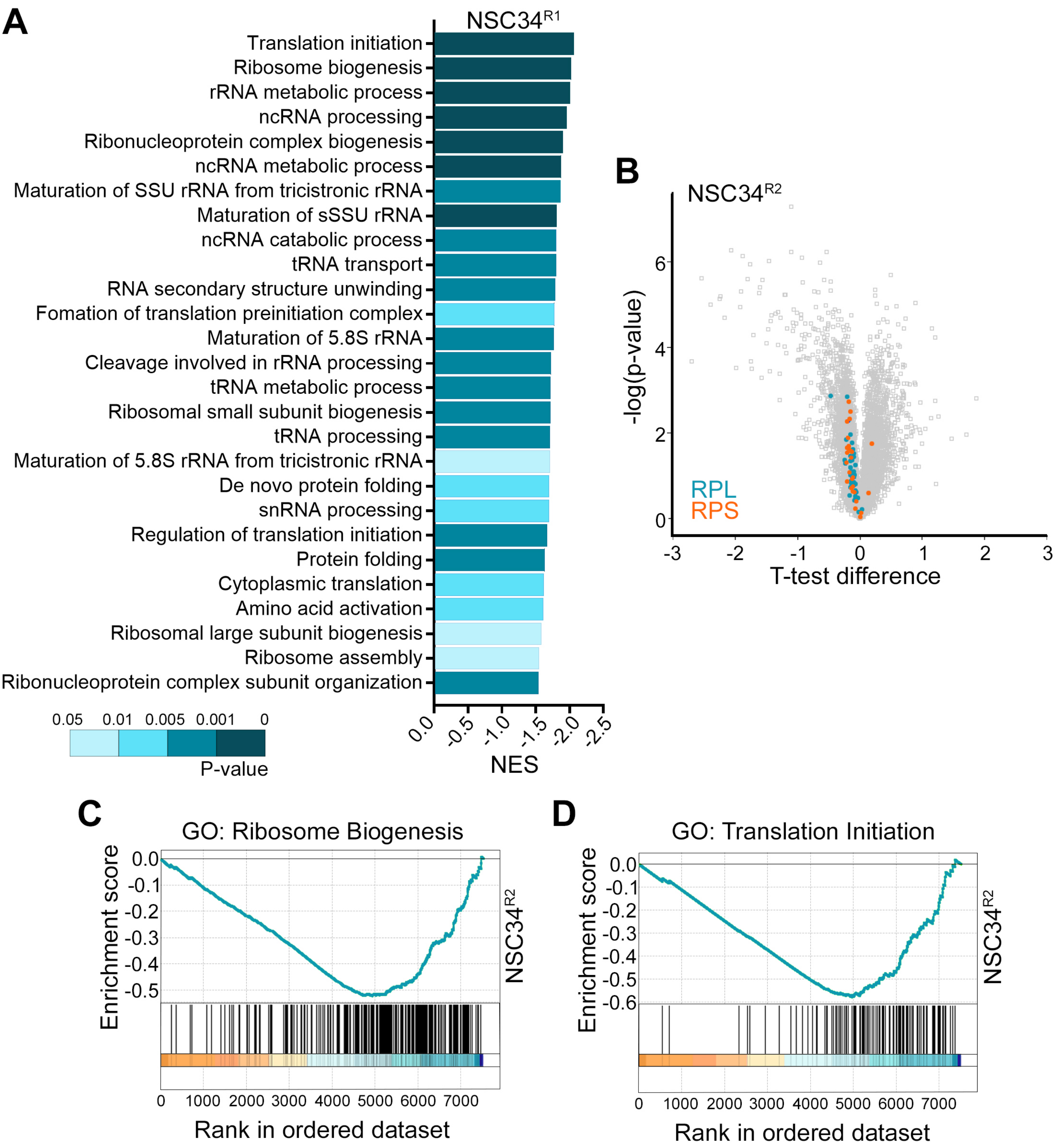
A
NSC34R1
Translation initiation
Ribosome biogenesis
rRNA metabolic process
ncRNA processing
Ribonucleoprotein complex biogenesis
ncRNA metabolic process
Maturation of SSU rRNA from tricistronic rRNA
Maturation of sSSU rRNA
ncRNA catabolic process
tRNA transport
RNA secondary structure unwinding
Fomation of translation preinitiation complex
Maturation of 5.8S rRNA
Cleavage involved in rRNA processing
tRNA metabolic process
Ribosomal small subunit biogenesis
tRNA processing
Maturation of 5.8S rRNA from tricistronic rRNA
De novo protein folding
snRNA processing
Regulation of translation initiation
Protein folding
Cytoplasmic translation
Amino acid activation
Ribosomal large subunit biogenesis
Ribosome assembly
Ribonucleoprotein complex subunit organization
0.0 -0.5 -1.0 -1.5 -2.0 -2.5
NES
0.05 0.01 0.005 0.001 0
P-value
B
NSC34R2
-log(p-value)
T-test difference
RPL
RPS
C
GO: Ribosome Biogenesis
Enrichment score
NSC34R2
Rank in ordered dataset
D
GO: Translation Initiation
Enrichment score
NSC34R2
Rank in ordered dataset

**Figure S5. PI3K, IGF1 or AKT inhibition trigger a transcriptional signature resembling to that of (PR)n-resistant cells.** (A, C) CMap connectivity scores (CS) of the PI3K (A), IGF1 (B) and (AKT) inhibitor compound sets, illustrating the similarity of their transcriptional signature to that of $NSC34^{R1}$ and $NSC34^{R2}$ cell. The CS is represented by the colour gradient, and the value indicated in each cell.

*Sirozh et al* **FigS5**

**A**

| PI3Ki | NSC34[R1] | NSC34[R2] |
|---|---|---|
| PIK-90 | 97.82 | 97.18 |
| AS-605240 | 97.35 | 98.50 |
| AZD-6482 | 96.55 | 71.55 |
| GDC-0941 | 96.51 | 97.60 |
| GSK-1059615 | 95.77 | 87.61 |
| PI-828 | 95.10 | 92.43 |
| ZSTK-474 | 91.92 | 87.02 |
| wortmannin | 87.91 | 94.67 |
| XL-147 | 54.10 | -81.87 |
| TGX-221 | 18.42 | 96.17 |
| AS-604850 | -84.44 | -49.60 |

**B**

| IGF1i | NSC34[R1] | NSC34[R2] |
|---|---|---|
| linsitinib | 97.82 | 91.79 |
| BMS-536924 | 92.75 | 95.32 |
| BMS-754807 | 92.57 | 94.33 |
| GSK-1904529A | 49.31 | 39.06 |
| PQ-401 | 20.84 | 22.19 |
| tyrphostin-AG-538 | -7.75 | 66.43 |
| I-OMe-AG-538 | -46.27 | 52.20 |
| EI-247 | -50.03 | -55.11 |

**C**

| AKTi | NSC34[R1] | NSC34[R2] |
|---|---|---|
| MK-2206 | 99.22 | 99.93 |
| AKT-inhibitor-1-2 | 99.08 | 99.33 |
| A-443644 | 97.82 | 64.53 |
| pyrvinium-pamoate | 91.83 | -65.22 |
| honokiol | 91.02 | -11.06 |
| AKT-inhibitor-IV | 28.54 | -35.19 |
| hexamethylenebisacetamide | 13.47 | 12.97 |
| triciribine | -55.64 | -27.62 |
| BML-257 | -60.78 | 36.25 |

**Figure S6. Characterization of the progeroid phenotype in $(PR)_{97}^{HA}$-expressing mice.** (A) Scheme representing the inducible double knock-in *Col1a1*$^{(PR)97}$ *Rosa26* $^{rtTA}$ *strategy* that enables dox-dependent expression of $(PR)_{97}^{HA}$ peptides in mice. (B) Table indicating the levels and subcellular localisation of $(PR)_{97}^{HA}$ expression, as evaluated by HA immunohistochemistry in different tissues in three representative $PR^{KI/KI}$ animals, 3 weeks after treatment with dox in the drinking water. (C) Evolution of body weights of non-inducible ($PR^{+/+}$) and inducible ($PR^{KI/KI}$) littermates after 0, 1 or 4 weeks of dox-treatment (0 and 1 weeks: $n^{PR+/+} = 12$, $n^{PR\ KI/KI} = 15$; 4 weeks: $n^{PR+/+} = 10$, $n^{PR\ KI/KI} = 10$) (n. s. $P > 0.05$; ****$P < 0.0001$; *t*-test). (D) Quantification of the skin thickness in non-inducible $PR^{+/+}$ ($n = 7$) and inducible $PR^{KI/+}$ ($n = 6$) littermates after 20 weeks of treatment with dox. Representative images are shown in **Fig. 4F**. (E) Representative CT images illustrating the onset of kyphosis in $PR^{KI/KI}$ mice after 12 weeks of treatment with dox. The kyphotic angle was quantified between the C1-T3-T9 vertebrae. (F) Quantification of the kyphotic angle from the data shown in (E).

A

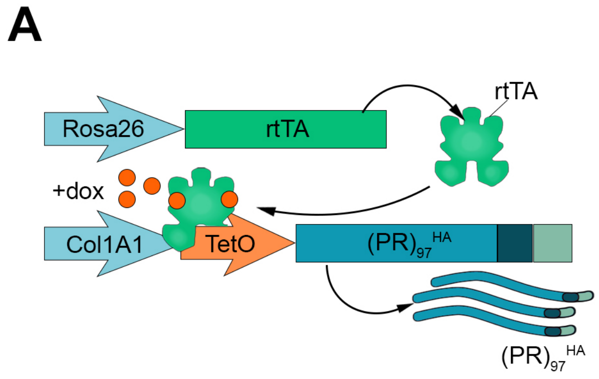


B

| Tissue | Expression | Localisation |
|---|---|---|
| Heart | + | N |
| Lungs | + | C, N, NO |
| Intestine | ++++ | C, N, NO |
| Liver | +++ | C, N, NO |
| Kidney | ++ | C, N, NO |
| Pancreas | ++ | C, N, NO |
| Thymus | +++ | C, N, NO |
| Bone Marrow | +++ | C, N, NO |
| Spleen | +++ | C, N, NO |
| Skin | ++ | C, N, NO |
| Brain | - | N/A |
| Spinal cord | - | N/A |

*C = cytoplasmic, N = nuclear, NO = nucleolar*

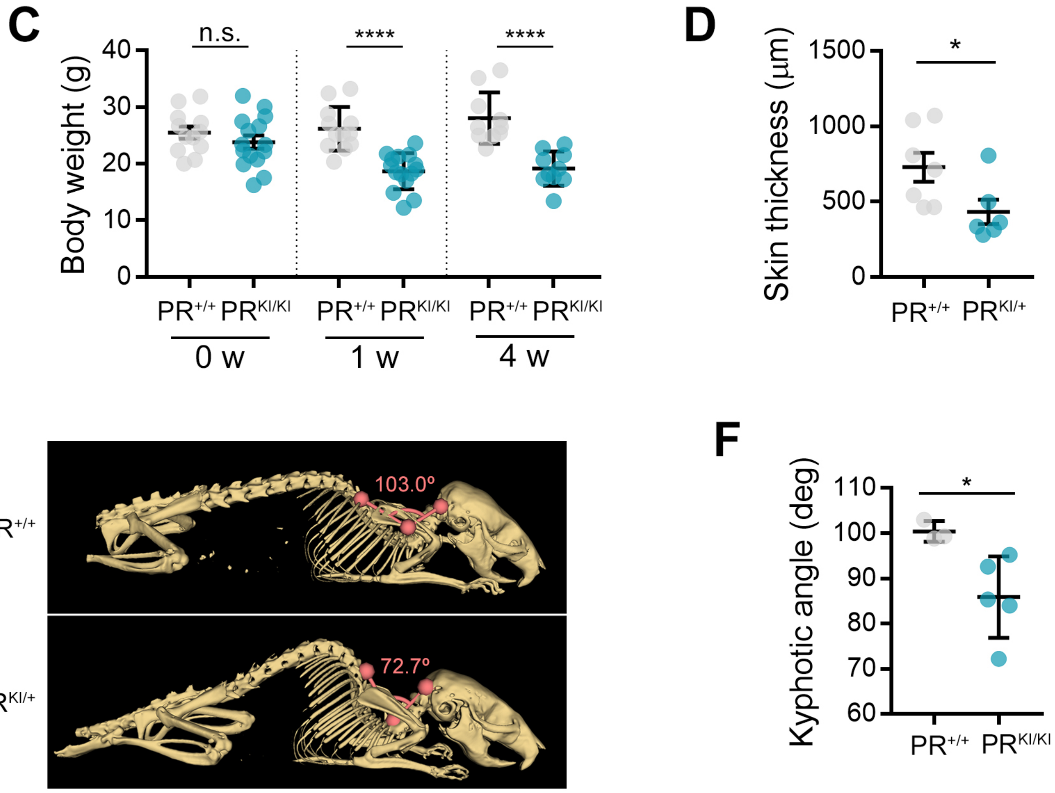

**Figure S7. NS and reduced translation upon expression in $(PR)_{97}^{HA}$ MEF.** (A) Representative IF of $(PR)_{97}^{HA}$ (red) and FBL (green) (A) in inducible $PR^{KI/KI}$ MEF 48 hrs after dox treatment. Scale bar represents 10 μm. (B) Average normalised viability, as evaluated with a CellTiter-Glo luminescent assay, of three non-inducible $PR^{+/+}$ MEF lines, and three $PR^{KI/KI}$ MEF lines, treated with dox for 48 hrs. Data information: n. s. $P > 0.05$; $^{*}P < 0.05$; $^{**}P < 0.01$; $^{****}P < 0.001$; paired *t*-test. (C) Representative image from cultures of $PR^{KI/KI}$ MEF grown in the presence or absence of dox for 48 hrs. (D, E) HTM-mediated quantification of HA intensity (D) and nucleolar area (E) per nucleus in $PR^{+/+}$ and $PR^{KI/KI}$ MEF, after 48 hrs of treatment with dox. (F) HTM-mediated quantification of HPG levels per cell in three non-inducible $PR^{+/+}$ MEF lines, and three inducible littermate $PR^{KI/KI}$ MEF lines, treated with dox for 48 hrs. HPG was added 1 hour prior to fixation. Black lines indicate mean values.

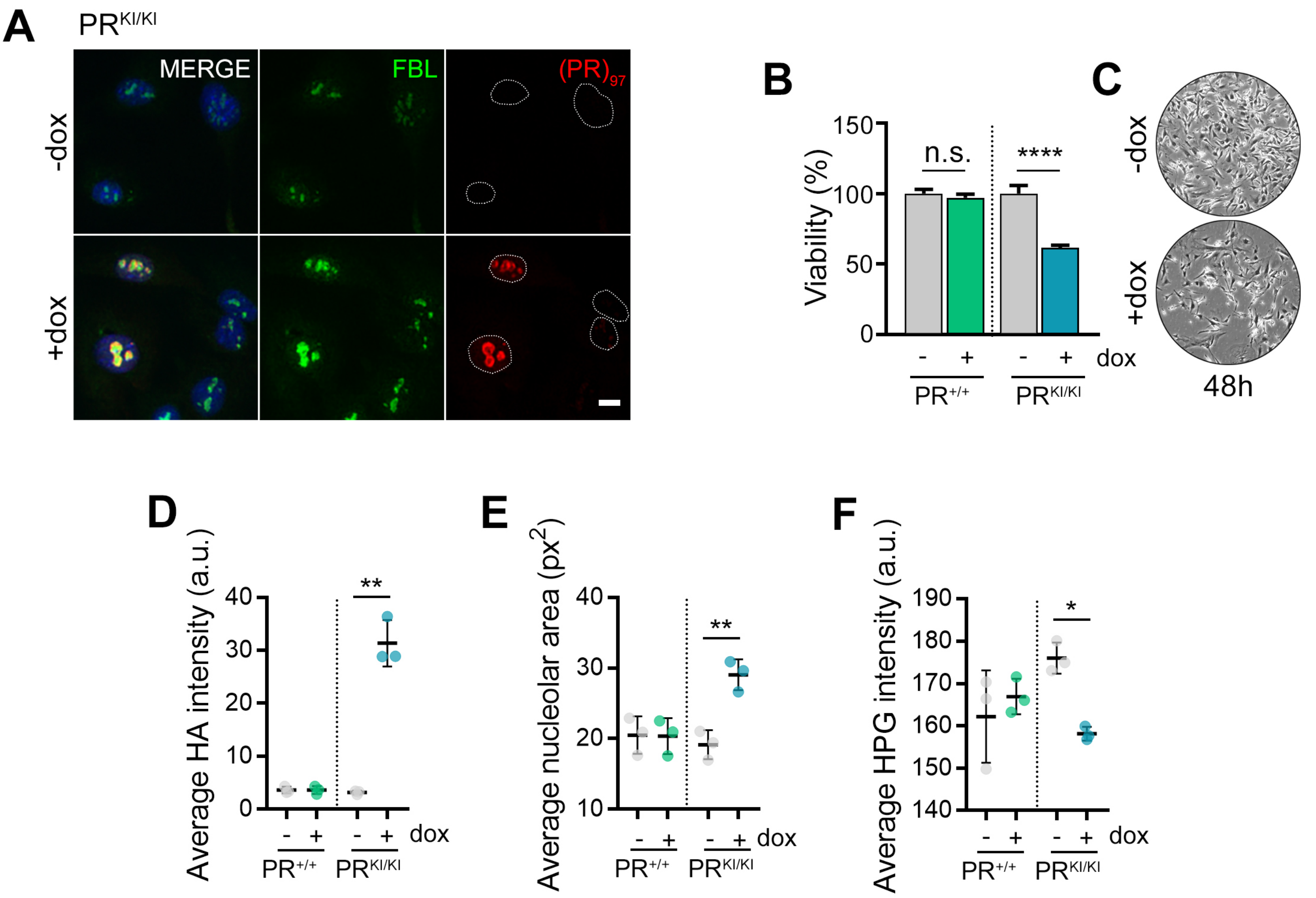
A
PR KI/KI
MERGE
FBL
(PR)97
-dox
+dox
B
Viability (%)
150
100
50
0
n.s.
****
- + - + dox
PR+/+
PR KI/KI
C
-dox
+dox
48h
D
Average HA intensity (a.u.)
40
30
20
10
0
**
- + - + dox
PR+/+
PR KI/KI
E
Average nucleolar area (px2)
40
30
20
10
**
- + - + dox
PR+/+
PR KI/KI
F
Average HPG intensity (a.u.)
190
180
170
160
150
140
*
- + - + dox
PR+/+
PR KI/KI

**Figure S8. mTOR inhibition rescues $(PR)_{97}^{HA}$-driven NS and toxicity MEF.** (A) IF of $(PR)_{97}^{HA}$ peptides (red) in intestine sections from littermate $PR^{+/+}$ and $PR^{KI/KI}$ mice, 3 weeks after treatment with dox in the drinking water. (B) Quantification of nucleolar FBL+ area from the images shown in (A). (C) IF of $(PR)_{97}^{HA}$ (red) and FBL (green), in $PR^{KI/KI}$ MEF after 48hrs of treatment with dox, in the presence or absence of rapamycin (1nM). Scale bar (white) represents 5 μm. (D, E) Quantification of the nucleolar area (D) and viability (E), as quantified by a CellTiter-Glo luminescent assay, in the experiment defined in (C). (n. s. $P > 0.05$; * $P < 0.05$; **** $P < 0.0001$; *t*-test).

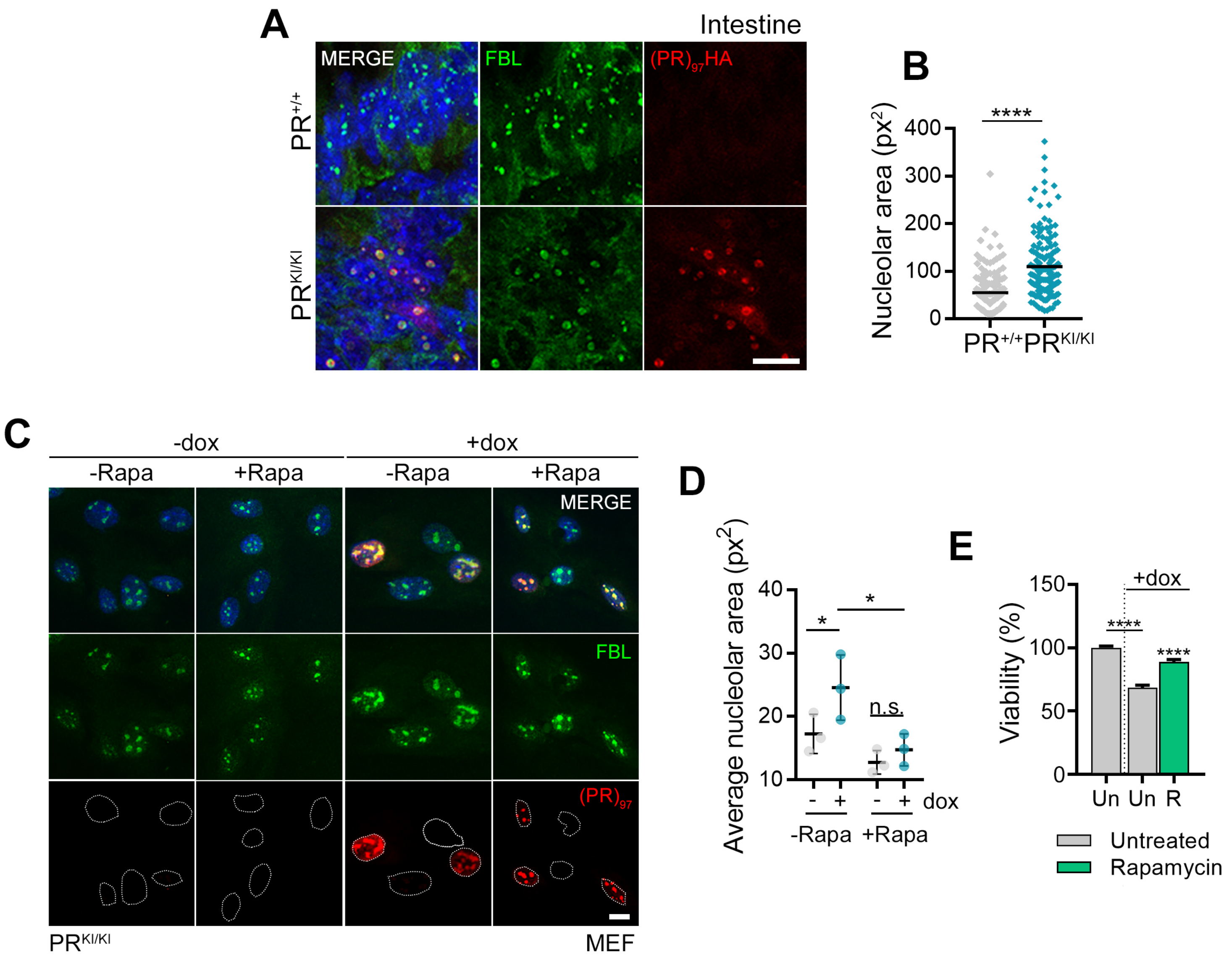
A
Intestine
MERGE
FBL
(PR)97HA
PR+/+
PRKI/KI
B
Nucleolar area (px2)
****
400
300
200
100
0
PR+/+ PRKI/KI
C
-dox
+dox
-Rapa
+Rapa
-Rapa
+Rapa
MERGE
FBL
(PR)97
PRKI/KI
MEF
D
Average nucleolar area (px2)
40
30
20
10
*
*
n.s.
- + - + dox
-Rapa +Rapa
E
Viability (%)
150
100
50
0
+dox
****
****
Un Un R
Untreated
Rapamycin

**Figure S9. Motorneurons derived from patients of *C9ORF72*-ALS present increased expression of r-genes and mTOR signalling factors.** (A, B) Volcano plot of RNA-seq data from differentiated motor neurons, derived from healthy controls (n=4), and patients of C9ORF72 ALS (n=8), available at the NeuroLINCS data repository. The grey points indicate gene expression distribution in all cases, while the coloured points correspond to genes defined as the “mTORC1 signalling” hallmark (A; green), and ribosomal proteins (B; blue and orange, RPLs and RPSs, respectively). Each volcano plot is accompanied by a bar chart summarising the gross distribution of each gene set in the corresponding expression profile analysis. The *P*-value was obtained by a hypergeometric test.

*Sirozh et al* **FigS9**

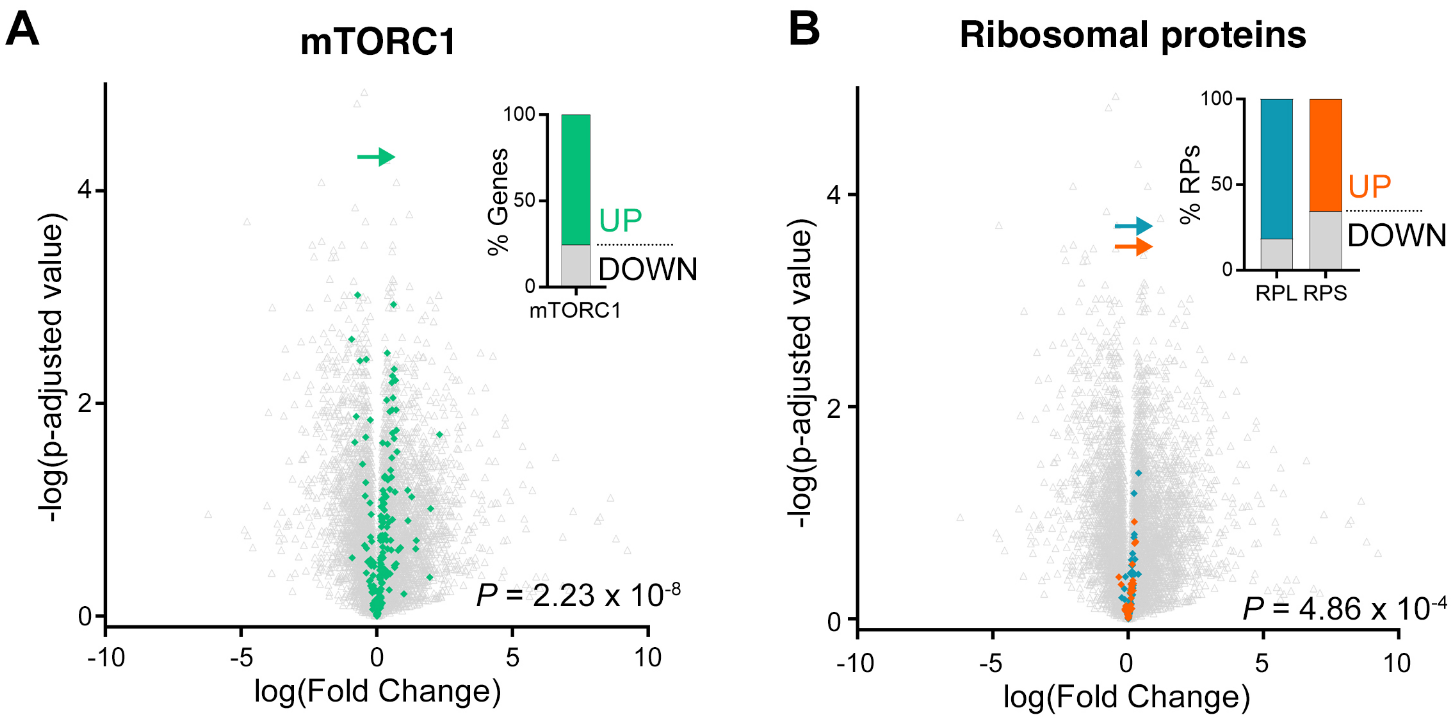


**Motoneuron RNAseq: healthy vs. C9ORF72 ALS**
